# Empirical optimization of dCas12a-based CRISPRi/a for pooled screening in *Yarrowia lipolytica*

**DOI:** 10.64898/2026.11.11.766072

**Authors:** Laetitia Maroc, Lukas Meyvisch, Penelope Smeets, Daan Troubleyn, Sibylle C. Vonesch

## Abstract

*Yarrowia lipolytica* is an important host for biotechnology and metabolic engineering, valued for its ability to accumulate lipids and produce a wide range of industrially relevant compounds. This creates a growing need for genome-scale tools that can accelerate the identification of genes underlying complex phenotypes. Although Cas9-based CRISPR interference (CRISPRi) and activation (CRISPRa) tools are well established in *Y. lipolytica*, Cas12a offers a complementary platform with a T-rich PAM that can expand access to promoter regions and facilitate multiplexed regulation. Despite these advantages, Cas12a-based CRISPRi/a remains comparatively underdeveloped in *Y. lipolytica,* particularly for large-scale functional screening. Here, we establish and systematically benchmark dCas12a-based CRISPRi/a platforms for pooled screening in *Y. lipolytica*. We first compared two widely used effectors, dLbCas12a and denAsCas12a, across single-gene assays and a pooled CRISPRi library targeting nine essential or conditionally essential genes with 752 crRNAs. Both orthologs supported robust transcriptional repression, but their relative performance was crRNA-and condition-dependent, with dLbCas12a showing more consistent and stronger late-stage depletion across targets. dLbCas12a also enabled enrichment-based CRISPRi and CRISPRa screens, recovering expected drug-resistance and lipid-associated phenotypes.

The broad design of the pooled library further enabled identification of crRNA features associated with CRISPRi efficacy. Highly active crRNAs were enriched near transcription start sites, with an effective targeting interval extending from approximately −260 to +50 bp. Intermediate GC content was associated with higher efficacy, while PAM identity showed a weaker but consistent effect, with TTTA and TTTT PAMs linked to stronger depletion than TTTC and TTTG PAMs. Spacer length showed the strongest association with CRISPRi efficacy, with 18-nt crRNAs consistently outperforming matched longer 20-, 23-, and 25-nt variants. Finally, we tested dHyperLbCas12a, an engineered LbCas12a variant with enhanced activity, and found that it produced substantially stronger CRISPRi and CRISPRa phenotypes than dLbCas12a. Together, these results establish Cas12a as a versatile platform for pooled CRISPRi/a screens in *Y. lipolytica*, identify practical crRNA prioritization rules for improved library performance, and highlight dHyperLbCas12a as a promising effector for future large-scale gain-and loss-of-function screens.

## Introduction

Non-conventional yeasts are increasingly exploited as microbial cell factories because they possess unique metabolic and physiological traits that are absent or less pronounced in traditional model organisms^1^. Their ability to utilize diverse feedstocks, tolerate industrially relevant conditions, and produce a wide range of valuable metabolites makes them attractive platforms for biotechnology and synthetic biology. Among these organisms, *Yarrowia lipolytic*a has emerged as one of the most extensively developed chassis for industrial biotechnology^2^. Its native oleaginous metabolism, ability to utilize a broad spectrum of carbon sources, and suitability for production of both lipid-derived and non-lipid products have motivated extensive development of genetic and synthetic biology tools for this organism^3,4^. As these tools have matured, an increasingly important challenge has shifted from the ability to engineer *Y. lipolytica* to identifying which genes should be perturbed, and in which direction, to obtain a desired phenotype.

Genome-scale CRISPR technologies have begun to address this challenge. Genome-wide Cas9 libraries have enabled identification of essential genes and engineering of lipid accumulation, morphology, substrate utilization and stress-tolerance phenotypes, while genome-wide measurements of Cas9 and Cas12a cutting activity have enabled identification of *Y. lipolytica*-specific gRNA design rules for effective gene disruption^5–7^. These approaches demonstrate the potential of pooled functional genomics in *Y. lipolytica*, but gene disruption captures only complete loss-of-function phenotypes, making transcriptional perturbation an important complement to knockout screening. CRISPR interference (CRISPRi) can create partial loss-of-function phenotypes, enabling interrogation of genes with a dosage-sensitive phenotype or for which complete disruption strongly compromises fitness. CRISPR activation (CRISPRa) enables gain-of-function phenotypes that cannot be recovered through knockout screening, which is particularly relevant for weakly expressed genes or cryptic functions: in *Y. lipolytica*, CRISPRa-mediated activation of the endogenous β-glucosidase-encoding genes *BGL1* and *BGL2* revealed a latent capacity for growth on cellobiose^8^.

CRISPR/Cas9-mediated transcriptional regulation is well-established in *Y. lipolytica.* A dCas9-Mxi1 CRISPRi platform was shown to efficiently repress endogenous genes, including transient suppression of the non-homologous end-joining machinery to increase homologous-recombination efficiency^9^, while a dCas9-VPR system enabled strong transcriptional activation of native genes^8,10^. More recently, a whole-genome CRISPRi library was used to identify genes associated with furfural and acetic-acid tolerance^11^. Cas12a provides an attractive complementary system to Cas9, including a T-rich PAM for targeting AT-rich yeast promoters, compact crRNA architecture, and intrinsic ability to process individual crRNA arrays from a single precursor array, which greatly facilitates multiplexing^12^. Despite these attractive properties, dCas12a-based CRISPRi/a screening platforms in *Y. lipolytica* remain underdeveloped compared with Cas9-based systems. Two questions are particularly important for extending Cas12a to pooled transcriptional perturbation screens. First, Cas12a ortholog choice can strongly influence activity across organisms, yet this has not been systematically evaluated in *Y. lipolytica*. Second, although crRNA position relative to the transcription start site (TSS) is a well-established determinant of CRISPRi/a activity^13–17^, the contribution of other crRNA features remains much less understood. Most studies of Cas12a-mediated CRISPRi/a in *Y. lipolytica* have focused on individual genes, reporters, or small crRNA sets^18^, leaving the broader determinants of activity largely unexplored. Notably, an early dCas12a study testing ten guides across a reporter found no clear relationship between target position and repression efficiency^19^. Addressing these questions directly in *Y. lipolytica* will therefore be important for development of efficient dCas12a-based screening platforms in this yeast.

Here, we establish a pooled dCas12a-based CRISPRi and CRISPRa framework for *Y. lipolytica*. We systematically compare two Cas12a orthologs, dLbCas12a and denAsCas12a, using a pooled CRISPRi library targeting essential and conditionally essential genes and show that both support robust repression while differing in the timing, magnitude, and crRNA dependence of their effects. Using dLbCas12a, we further demonstrate CRISPRi- and CRISPRa-based enrichment screens across growth-based drug selections and FACS-based lipid-associated phenotypes. By deliberately sampling a broad targeting space, we empirically define a preferred TSS-proximal interval and identify sequence and spacer-length features associated with stronger CRISPRi-mediated depletion. Finally, we evaluate the engineered dHyperLbCas12a variant and show further enhancement of both repression and activation phenotypes. Together, these results provide a framework for Cas12a effector selection and crRNA prioritization and support the development of larger-scale gain- and loss-of-function screens in *Y. lipolytica*.

## Results

### Establishment of Cas12a-based CRISPRi and CRISPRa systems in *Y. lipolytica*

To establish a Cas12a-based transcriptional regulation platform for *Y. lipolytica*, we compared two nuclease-deficient Cas12a orthologs: dLbCas12a, derived from *Lachnospiraceae bacterium* ND2006, and denAsCas12a, derived from the engineered enAsCas12a variant, an optimized version of AsCas12a^20^. Each nuclease was fused to either the repressor Mxi1 or the activator VPR to generate CRISPRi and CRISPRa systems, respectively (Fig. 1A). To facilitate future pooled screening applications, all dCas12a effector cassettes were stably integrated at the neutral IntC_2 locus^21^, while crRNAs were expressed from a low-copy replicative plasmid (Fig. 1A). Prior to evaluating transcriptional regulation, we tested whether constitutive expression of the CRISPRi/a machinery affected host fitness. Growth of strains expressing dLbCas12a-Mxi1, dLbCas12a-VPR, denAsCas12a-Mxi1 or denAsCas12a-VPR was comparable to that of the parental W29 strain, indicating that all four dCas12a effector systems were well tolerated in *Y. lipolytica* (Fig. 1B).

**Figure 1.**
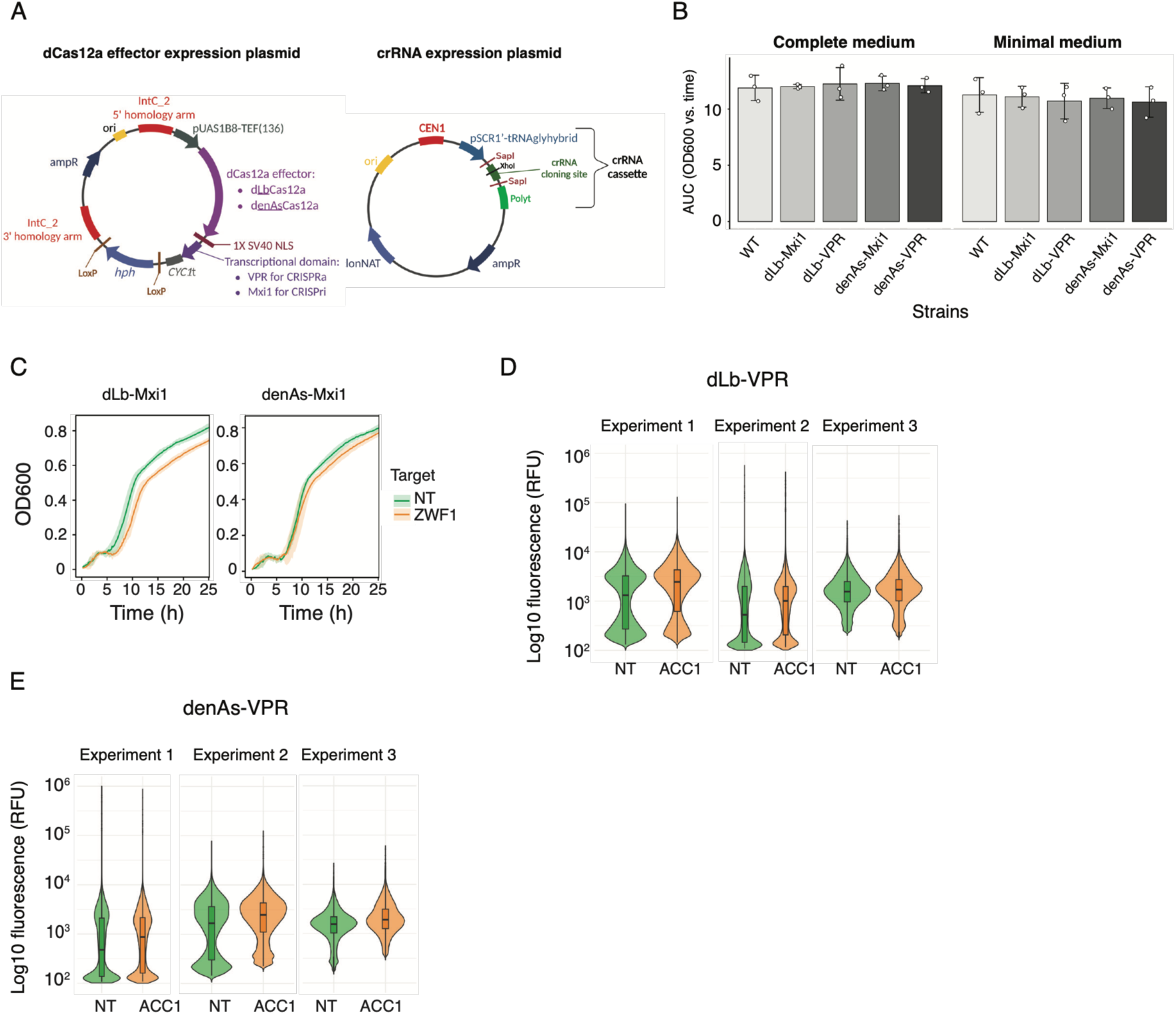
Establishment and initial testing of Cas12a-based CRISPRi and CRISPRa systems in *Y. lipolytica*. **(A)** Schematic representation of the CRISPRi/a systems. dCas12a effector cassettes (dLb-VPR, dLb-Mxi1, denAs-Mxi1 or denAs-VPR) were integrated at the neutral IntC_2 locus and expressed from the constitutive pUAS1B8-TEF(136) promoter. crRNAs were expressed from the constitutive SCR1’-tRNAGly promoter on a low-copy replicative plasmid carrying a Cas12a-specific direct repeat (LbDR or AsDR). **(B)** Expression of dCas12a effector proteins does not impair host fitness. Strains expressing dLbCas12a-Mxi1, dLbCas12a-VPR, denAsCas12a-Mxi1, or denAsCas12a-VPR were cultivated in complete or minimal medium and growth was monitored by OD600 measurements over 24 h. The relative area under the time-versus-OD curve (AUC) was then calculated. Dots represent the mean AUC, and error bars represent the standard deviation. **(C)** CRISPRi-mediated repression of *ZWF1*. Growth curves of strains expressing non-targeting (NT) or ZWF1-targeting crRNAs in dLbCas12a-Mxi1 and denAsCas12a-Mxi1 backgrounds. Experiments were performed independently two times. Within each experiment, four biological replicates were analyzed,each consisting of three technical replicates. Technical replicates were averaged prior to analysis, and the plotted values represent the mean of the biological replicates across the two independent experiments. Shaded ribbons indicate ±1 standard deviation. **(D,E)** CRISPRa-mediated transcriptional activation of *ACC1* using dLb-VPR (D) or denAs-VPR (E). Intracellular lipid accumulation was quantified by Nile Red staining and flow cytometry in strains expressing non-targeting (NT) or *ACC1*-targeting crRNAs. Violin plots show single-cell fluorescence distributions after exclusion of events overlapping with the corresponding unstained controls. The central line indicates the median fluorescence and the box represents the interquartile range. Each experiment consisted of four biological replicates performed on the same day, and cells from biological replicates were pooled within each experiment for visualization. Data shown are representative of three independent experiments.

We benchmarked both dCas12a orthologs using crRNAs targeting genes whose transcriptional perturbation produces easily measurable phenotypes. Repression of *ZWF1*, which encodes glucose-6-phosphate dehydrogenase, the first enzyme of the oxidative pentose phosphate pathway, was expected to reduce growth, providing a sensitive CRISPRi readout. Conversely, activation of *ACC1*, which encodes acetyl-CoA carboxylase, the enzyme catalyzing the first committed step in fatty acid biosynthesis, was expected to increase lipid accumulation. Changes in intracellular lipid content can be readily quantified by Nile Red staining coupled to flow cytometry, thereby providing a robust and sensitive CRISPRa phenotype. Together, these targets enabled a direct comparison of the CRISPRi/a performance of the two Cas12a orthologs in *Y. lipolytica*. Both dCas12a variants generated measurable CRISPRi and CRISPRa phenotypes, demonstrating functional transcriptional regulation with all effectors. For CRISPRi, repression of *ZWF1* produced a markedly stronger growth defect with dLbCas12a-Mxi1 than with denAsCas12a-Mxi1 (Fig. 1C). Conversely, activation of *ACC1* increased lipid accumulation to a similar extent with both effectors (Fig. 1D). Because these initial comparisons were based on only a single target gene for each application, they cannot establish whether the observed differences reflect general performance differences between orthologs or target-specific effects.

### Comparative benchmarking reveals stronger and more consistent CRISPRi activity with dLbCas12a in *Y. lipolytica*

To systematically compare the dLbCas12a-and denAsCas12a-based CRISPRi platforms across diverse targets, we constructed a small library targeting 13 endogenous genes (Table S1). The target set encompassed essential and conditionally essential functions, biosynthetic pathways and drug resistance, allowing CRISPRi activity to be assessed across diverse cellular processes and growth conditions. crRNAs were designed across a deliberately broad targeting region, extending from 1 kb upstream of the start codon to a position corresponding to 10 % of the annotated gene length downstream of the start codon, irrespective of exon-intron structure. This permissive design was chosen to empirically define a preferred CRISPRi targeting window and other CRISPRi efficacy associated features in *Y. lipolytica*. In addition to 23-nt crRNAs, matched 18-, 20-, 23- and 25-nt variants were included for a subset of targets to assess the effect of spacer length. The final library comprised 752 crRNAs, including 32 non-targeting controls (4.3 % of the library).

We first benchmarked CRISPRi performance between dLbCas12a-Mxi1 and denAsCas12a-Mxi1 by growing the pooled libraries in complete (SCC) and minimal (SCM) media and monitoring crRNAs targeting essential genes (6 genes, 300 crRNAs) and conditionally essential genes (3 genes, 159 crRNAs) genes. crRNA abundance was quantified over successive generations, as a measure of CRISPRi-associated fitness effects in each condition (Fig. 2A). Measurements were highly reproducible between biological replicates (ρ > 0.95; Fig. S1). At the experimental T0, crRNAs targeting essential genes were already depleted compared to non-targeting controls (Fig. 2B), consistent with constitutive repression causing fitness defects before the start of the experiment. The magnitude of this initial depletion was broadly comparable between the two Cas orthologs, indicating that both systems were active.

**Figure 2.**
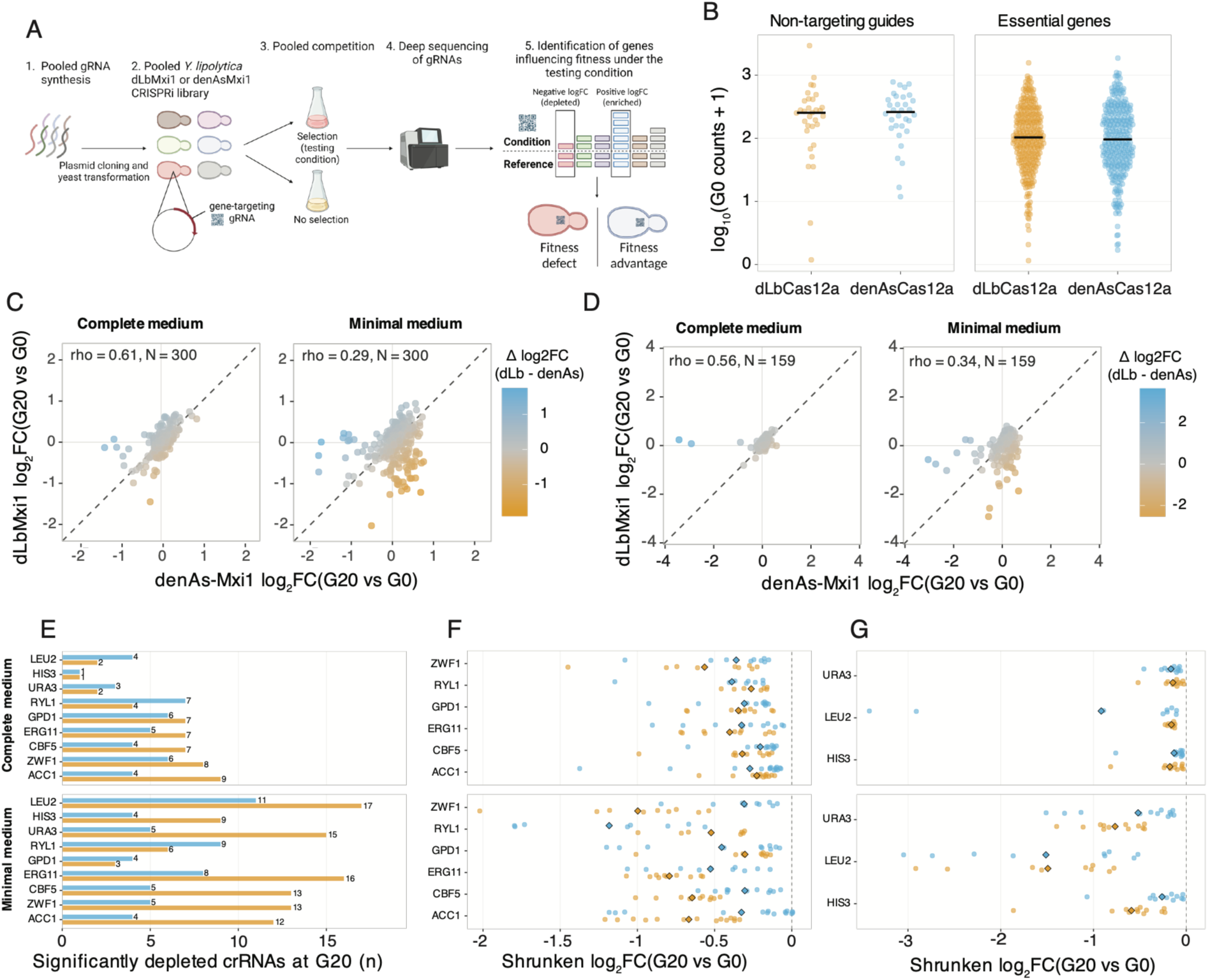
Comparative benchmarking of dLbCas12a-and denAsCas12a-based CRISPRi. **(A)** Experimental design of the pooled CRISPRi screen comparing dLbCas12a-Mxi1 and denAsCas12a-Mxi1 across growth conditions and generations. A pooled crRNA library was synthesized, cloned into a plasmid, and transformed into *Y. lipolytica* strains expressing either dLbCas12a-Mxi1 or denAsCas12a-Mxi1. The resulting pooled populations were propagated under selective and non-selective growth conditions, and crRNA abundances were quantified by NGS at successive generations. Changes in crRNA representation were then used to identify crRNAs associated with fitness defects or fitness advantages under the tested conditions. **(B)** Normalized crRNA abundance at G0 for non-targeting crRNAs and crRNAs targeting essential genes for both Cas variants. Points represent individual crRNAs; black bars indicate medians.**(C)** Comparison of matched crRNA-level log_2_ fold-changes (log_2_FCs) between denAsCas12a-Mxi1 and dLbCas12a-Mxi1 for essential genes (N = 300 crRNAs) at G20 (corresponding to 20 generations of competitive growth) in complete and minimal medium. Each point represents the same crRNA tested with both Cas variants; the dashed diagonal indicates equal effects between variants. **(D)** Matched crRNA-level comparison as in (C) for conditionally essential genes at G20 (N = 159 crRNAs). **(E)** Number of significantly depleted (padj <= 0.05) crRNAs per target gene at G20 for dLbCas12a-Mxi1 and denAsCas12a-Mxi1 under the relevant growth conditions. **(F–G)** Comparison of CRISPRi effect magnitude among the most depleted 20% (based on log_2_FC) of crRNAs for each target gene, selected independently within each Cas variant and condition, at G20. **(F)** essential genes and **(G)** conditionally genes. More negative log_2_FC values indicate stronger crRNA-associated fitness defects. Diamond indicates mean of 20% most depleted crRNAs per gene and Cas variant. In B, E-G, gold = dLbCas12a-Mxi1, light blue = denAsCas12a-Mxi1.

We next compared crRNA-level depletion over subsequent generation. Both Cas12a variants supported functional CRISPRi, with crRNAs targeting essential and conditionally essential genes becoming progressively depleted during growth. However, matched-crRNA comparisons revealed substantial differences in both depletion magnitude and the identity of the most strongly depleted crRNAs between the two orthologs (Fig 2C; Fig. S2A). Log_2_-fold changes (log_2_FCs) for crRNAs targeting essential genes in complete medium were moderately concordant between dLbCas12a-Mxi1and denAsCas12a-Mxi1 (Spearman ρ = 0.61-0.64 after 15 -25 generations), whereas concordance was lower in minimal medium (ρ = 0.29-0.31; Fig. 2C; Fig. S2A). A similar reduction in concordance was observed for conditionally essential genes (Fig 2D, Fig. S2B). In contrast, non-targeting controls showed substantially higher concordance between Cas orthologs (Fig. S2C), indicating that observed differences were primarily crRNA-dependent rather than reflecting general experimental variability.

The relative performance of the two orthologs also changed over time. In complete medium, where depletion was expected primarily for essential gene targeting crRNAs, dLbCas12a-Mxi1 generally produced an equal or greater number of significantly depleted crRNAs per gene (Fig. 2E; Fig. S3A). In minimal medium, denAsCas12a-Mxi1 showed evidence of earlier activity, whereas dLbCas12a-Mxi1 generally produced more significantly depleted crRNAs at later generations and across both media (Fig. 2E; Fig. S3A). To determine whether dLbCas12a-Mxi also produced stronger phenotypes among relatively active crRNAs, we compared the log_2_FCs of the 20% most depleted crRNAs within each gene, condition and Cas ortholog. Because the library was intentionally designed to sample a broad targeting space, many crRNAs were expected to show little or no activity. Focusing on the most depleted guides therefore allowed us to compare the stronger-performing fraction without diluting their effects through whole-gene averages. dLbCas12a-Mxi1 showed a more negative mean log_2_FC for five of six essential genes and two of three conditionally essential genes at later generations (G20, G25). Conversely, at G10 and G15, denAsCas12a-Mxi1 generally showed more negative mean log2FCs, consistent with its higher number of significantly depleted crRNAs at these earlier time points, although the effects were generally modest (Fig. 2F,G; Fig. S3B,C). Together, these results indicate that denAsCas12a-Mxi1 can exhibit faster initial activity, whereas dLbCas12a-Mxi1 provides stronger and more consistent repression at later generations across genes and growth conditions. Based on this broader and more consistent performance, we selected dLbCas12a-Mxi1 for subsequent experiments.

### dLbCas12a CRISPRi/a enables robust detection of enrichment phenotypes

We next asked whether dLbCas12a could also identify crRNA-associated enrichment phenotypes, complementing the depletion-based screens described above. As a CRISPRi test case, we targeted *CAN1*, which encodes an arginine permease, during growth in L-canavanine. L-canavanine is a toxic arginine analog, and reduced *CAN1* expression is therefore expected to enrich for efficient *CAN1*-targeting crRNAs. Accordingly, we observed strong enrichment of multiple *CAN1*-targeting crRNAs, with effects most pronounced after 10-20 generations of competitive growth (Fig. 3A, Fig. S4).

**Figure 3.**
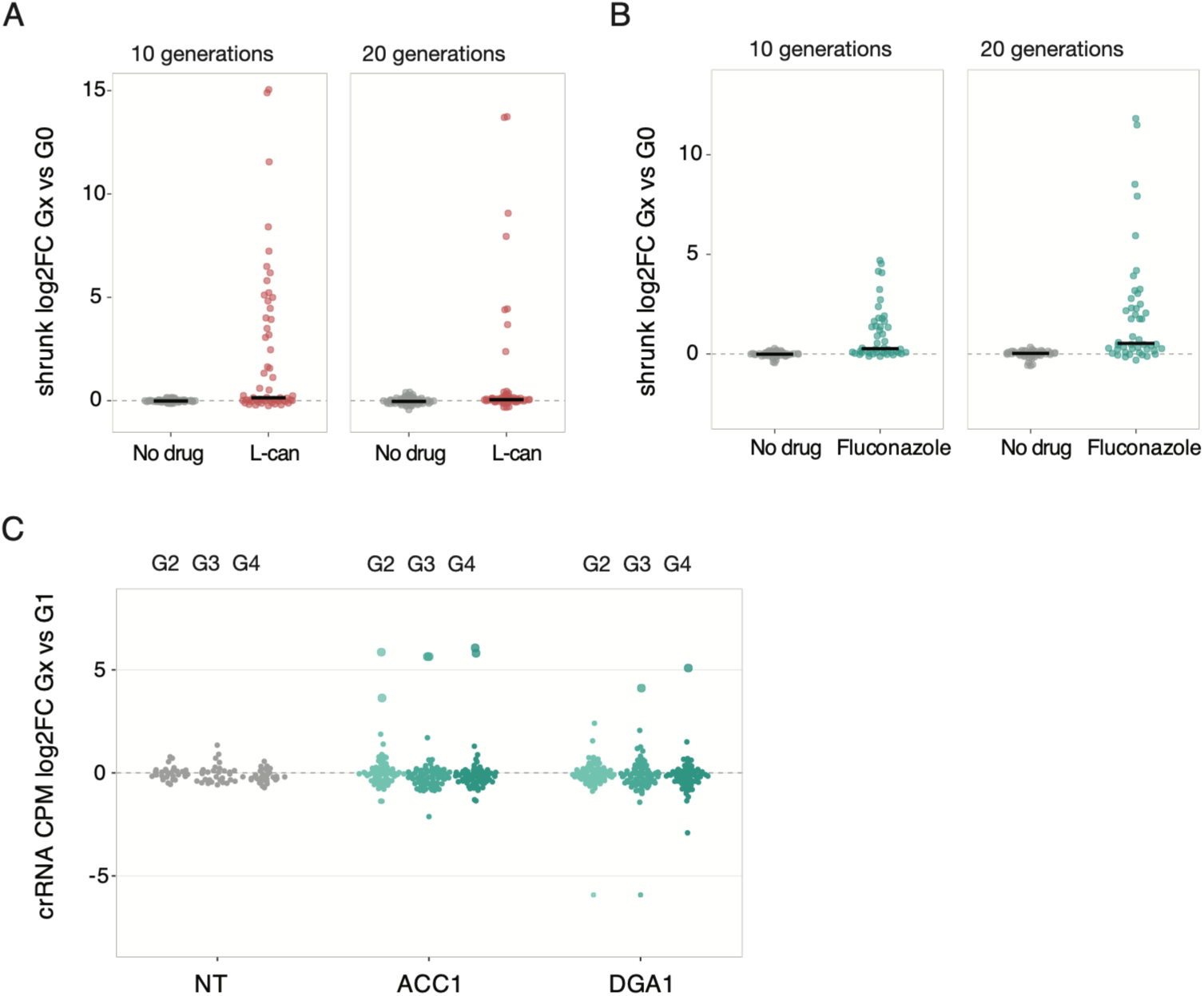
dLbCas12a supports CRISPRi- and CRISPRa-based phenotypic enrichment screens. **(A)** CRISPRi targeting of *CAN1* during growth in medium with (red) or without (grey) L-canavanine. CrRNA-level log_2_fold-changes (log_2_FCs) are shown across generations. Enrichment of *CAN1*-targeting crRNAs under L-canavanine selection is consistent with reduced *CAN1* expression conferring resistance. **(B)** CRISPRa targeting of *ERG11* during growth in complete medium with (turquoise) or without (grey) fluconazole. CrRNA-level log_2_FCs are shown across generations. Enrichment of *ERG11*-targeting crRNAs under fluconazole selection is consistent with increased *ERG11* expression promoting resistance. **(C)** CRISPRa screen for altered lipid accumulation following targeting of *ACC1* and *DGA1*. Cells were stained with Nile Red and sorted by FACS into four gates spanning baseline to increased lipid-associated fluorescence (G1 < G2 < G3 < G4). CrRNA enrichment relative to the baseline G1 population is shown for replicate 2 for G2-G4 (left to right) for nontargeting (grey), and *ACC1*-/*DGA1*-targeting crRNAs (in turquoise shades). Positive log2FC values indicate enrichment relative to G1. Larger points indicate crRNAs with zero counts in the G1 reference gate and non-zero counts in the corresponding higher-lipid gate.

We then tested CRISPRa in two distinct applications. First, we targeted *ERG11*, which encodes the enzyme lanosterol 14-alpha-demethylase, the target of fluconazole, reasoning that higher *ERG11* expression could increase fluconazole resistance. Consistent with this, *ERG11* activation produced a clear enrichment phenotype under fluconazole selection, with multiple *ERG11*-targeting crRNAs becoming progressively enriched over time (Fig. 3B, Fig. S4). In addition to demonstrating robust dLbCas12a-VPR CRISPRa activity in *Y. lipolytica*, these results present direct evidence that increased *ERG11* expression can enhance fluconazole tolerance in *Y. lipolytica*.

As a second CRISPRa application, we investigated a metabolic phenotype by targeting *ACC1* and *DGA1*, two key lipid biosynthesis genes whose increased activity is expected to promote lipid accumulation. Cellular lipid content was quantified by Nile Red staining followed by FACS, with cells sorted into four gates spanning baseline (G1), medium (G2), high (G3), and very high (G4) lipid-associated fluorescence. crRNA abundances were highly concordant between biological replicates across gates (Spearman ρ = 0.89-0.95 for G1-G3, Fig. S5A). Concordance was somewhat lower for the highest lipid gate (G4), for which replicate 1 had substantially lower sequencing depth and crRNA coverage (15x versus >67x in other samples). Several *ACC1*-and *DGA1*-targeting crRNAs showed greater enrichment between G2-4 relative to G1 than non-targeting controls (Fig. 3C, Fig. S5), although the FACS phenotype was more noisy and generally fewer effective crRNAs were identified than for the growth or drug-resistance screens. To identify reproducible FACS phenotypes, we required crRNAs to show a log2FC > 1 in both biological replicates within the same gate. By this criterion, *ACC1* emerged as the most consistent hit, with enriched crRNAs detected across all three higher fluorescence gates (G2-G4), whereas *DGA1* enrichment was restricted to G3 and G4. Because overexpression of the essential gene *ZWF1* has previously been shown to enhance lipid accumulation in *Y. lipolytica*^22^, we also examined whether crRNAs targeting *ZWF1* or other genes were enriched in the higher fluorescence gates. crRNAs targeting *ZWF1* were reproducibly enriched in G2 and G3, while *CBF5*-targeting crRNAs were enriched in all high-fluorescence gates. Notably, no non-targeting crRNAs met this enrichment threshold, supporting the specificity of these signals. The recovery of *ZWF1* is consistent with its previously reported effect on lipid accumulation. In contrast, *CBF5* has not, to our knowledge, been linked to this phenotype. Because Nile Red fluorescence may reflect cellular properties other than simply lipid abundance, the *CBF5* phenotype should be interpreted cautiously and may reflect an indirect effect rather than increased lipid accumulation. Together, these experiments show that dLbCas12a supports both CRISPRi- and CRISPRa-based enrichment screens across distinct phenotypic readouts.

### Empirical determinants of CRISPRi efficacy

crRNA position relative to the TSS is a well-established determinant of CRISPRi efficacy, although the optimal targeting window can differ between effectors and species^23^. To identify the positional preference in *Y. lipolytica*, we focused on log_2_FCs of crRNAs targeting essential and conditionally essential genes in minimal medium (SCM) at G20, which provided the largest informative crRNA set and effects were well established by this timepoint. This analysis revealed that the most strongly depleted crRNAs clustered in a relatively narrow region around and upstream of the TSS (Fig. S6A), broadly consistent with the promoter-proximal targeting preference reported for CRISPRi in other organisms^15,24,25^. A comparable promoter-proximal pattern was observed for *ERG11* CRISPRa, although significantly enriched crRNAs extended over a broader region upstream of the TSS (Fig. S6B).

To quantify the relationship between TSS distance and CRISPRi activity, we performed a sliding-window analysis of gene-centered crRNA efficacy, defined as −log_2_FC such that stronger depletion corresponded to higher efficacy. Because our deliberately permissive crRNA-design strategy generated a library containing both active and largely inactive crRNAs, we focused on the upper tail of crRNA efficacy rather than average performance within each window, using the 80th percentile as a cutoff for relatively high-performing crRNAs. The most broadly supported increase in efficacy occurred immediately upstream of the TSS, with windows centered at −75, −50, −25 and 0 bp, each represented by at least seven of the nine genes included in the analysis, consistently showing elevated efficacy (Fig. 4A). A second peak was observed at −250 to −225 bp, although this region was supported by only five genes. Together, these data indicate that crRNAs mediating the strongest CRISPRi depletion in *Y. lipolytica* are preferentially concentrated at or immediately upstream of the TSS, while active crRNAs can occur across a broader promoter-proximal region. We therefore retained a −260 to +50 bp targeting window for subsequent analyses of sequence features associated with crRNA efficacy, capturing both the broadly supported TSS-proximal peak and highly active crRNAs observed further upstream (for e.g. *ACC1*) and immediately downstream of the TSS (for e.g. *RYL1, ZWF1, GPD1, URA3*).

**Figure 4.**
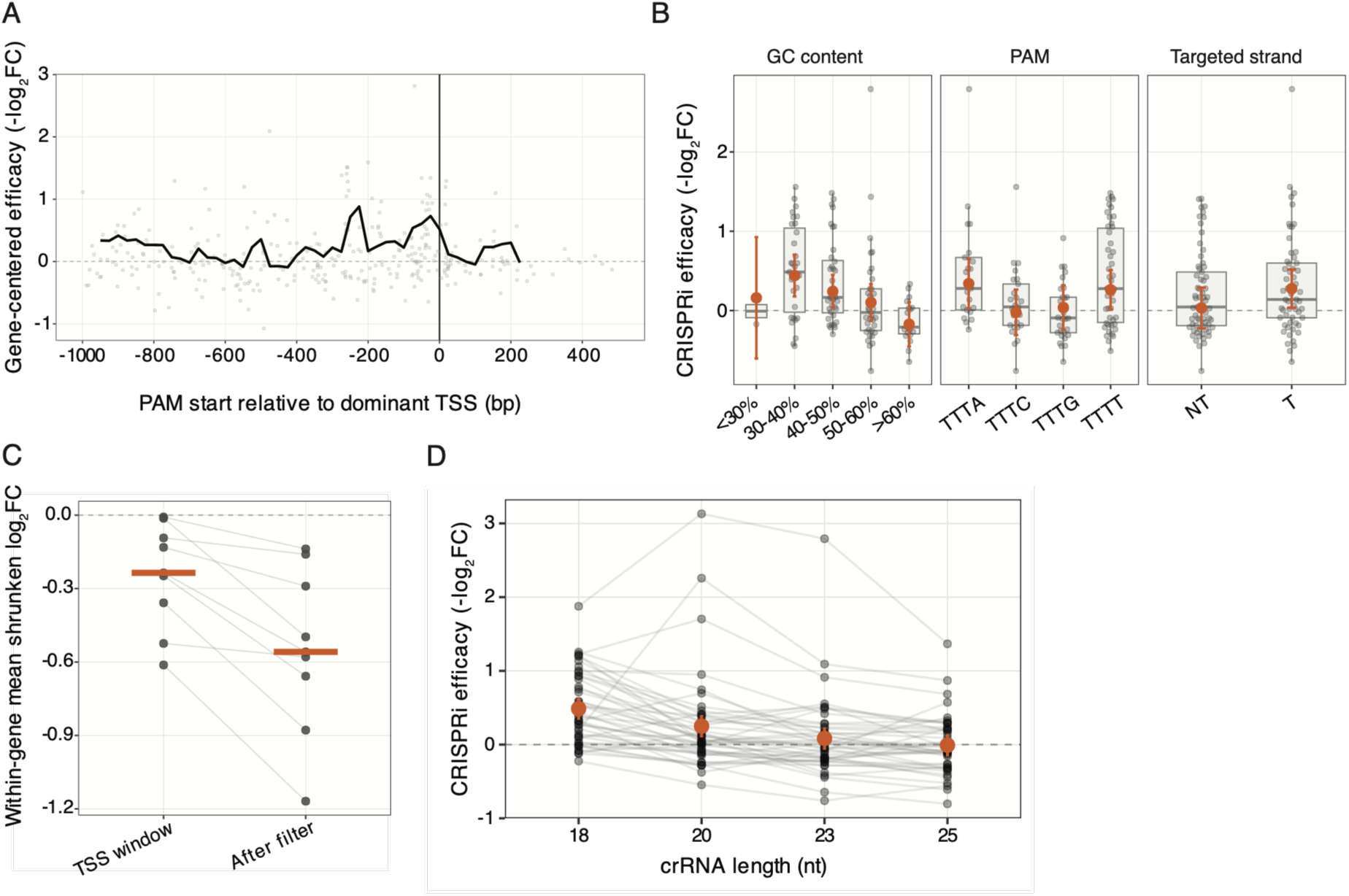
Empirical determinants of CRISPRi crRNA efficacy in *Y. lipolytica*. **(A)** Relationship between crRNA position relative to the dominant transcription start site (TSS) and CRISPRi efficacy. Grey points represent individual gene-centered crRNA efficacies. The black line represents the median of gene-specific 80th-percentile efficacies calculated across overlapping 100-bp windows at 25-bp intervals. **(B)** Associations between crRNA sequence features and CRISPRi efficacy within the −260 to +50 bp targeting window. Shown are GC-content class, PAM sequence and targeted strand. Individual crRNAs are shown as grey dots and boxplots, adjusted estimated marginal means with 95% confidence intervals from a linear model accounting for target gene, residual TSS-position effects and the other crRNA features are shown in orange. **(C)** Effect of applying the candidate sequence filter within the preferred TSS-targeting window. Mean crRNA log_2_FCwas calculated for each target gene, including only crRNAs in the predefine doptimal -260 to +50 bp window relative to the TSS (TSS window) and further removing any crRNAs with a GC content <30% or >60% and with a TTTC or TTTG PAM (after filtering). Lines connect values for the same gene. **(D)** Effect of spacer length on CRISPRi efficacy. Analysis was restricted to target sites represented by complete matched N18, N20, N23 and N25 crRNA sets, allowing spacer lengths targeting the same genomic position to be compared directly. Spacer-length effects were assessed using a linear mixed-effects model with target gene as a fixed effect and target site as a random intercept. Individual crRNAs are shown as grey points, with lines connecting spacer-length variants targeting the same site; orange points and error bars indicate adjusted estimated marginal means and 95% confidence intervals from the mixed-effects model.

Having defined a preferred targeting interval, we next asked whether additional crRNA features were associated with efficacy. We examined several features previously implicated in Cas12a activity in *S. cerevisiae* and mammalian systems^15,24,26,27^ including PAM identity, the base immediately upstream of the PAM, targeted strand and perturbation of the direct-repeat (DR) structure. We used a combined model that accounted for target gene and residual TSS-distance effects while testing each crRNA feature after accounting for the others. GC content was initially analyzed in bins to facilitate interpretation. GC content showed the clearest association with CRISPRi efficacy (partial F-test, *p* = 0.019, adjusted *p* = 0.096), with higher efficacy observed for 30–50% GC crRNAs compared to 50-60% GC, and a clear drop for >60% GC crRNAs (Fig. 4B). To exclude that this association resulted from the chosen GC-bin boundaries, we repeated the analysis with GC modeled as a continuous spline, to better represent its continuous nature. This independently supported an association between GC content and efficacy (*p* = 0.0019, adjusted *p* = 0.0093). PAM sequence and targeted strand showed weak effects (*p* = 0.059 and *p* = 0.058, respectively, Fig. 4B), but neither remained supported after multiple-testing correction. Adjusted efficacy was higher for TTTA and TTTT PAMs than for TTTG and TTTC PAMs, although no individual PAM comparison was significant. We found no evidence that either the nucleotide immediately upstream of the PAM (*p* = 0.499) or predicted DR perturbation (*p* = 0.920) was associated with CRISPRi efficacy in this dataset (Fig. S7A). Feature-associated differences became progressively clearer as depletion developed over successive generations. GC-content differences were minimal at G5– G10, began to emerge by G15 (*p* = 0.079), and were strongest at G20 and G25 (*p* = 0.019 and 0.032, respectively; Fig. S7B). PAM and targeted strand effects showed a similar consistent tendency. The progressive emergence of these feature-associated patterns alongside increasing CRISPRi depletion supports their biological relevance and argues against their being specific to the G20 time point. Finally, we asked whether CHOPCHOP v3 rank or predicted efficacy^28^ were associated with CRISPRi efficacy in *Y. lipolytica*. Although CHOPCHOP was initially developed for cleavage crRNA design, the most recent version (v3) also incorporates crRNA selection for CRISPRi. Relationships varied among target genes and showed no consistent trend (Fig. S8). Accordingly, after accounting for differences among target genes, predicted efficacy was not associated with CRISPRi efficacy (*p* = 0.163). Rank showed a weak trend (*p* = 0.064), but the effect was not in the expected direction, with higher (worse) CHOPCHOP ranks tending to be associated with stronger measured CRISPRi efficacy (β = 0.0185).

Collectively, these analyses identified GC content as the most strongly supported sequence-associated feature, with maximal adjusted efficacy at intermediate GC content and reduced efficacy at high GC content. PAM identity showed a weaker but consistent tendency, with TTTA and TTTT crRNAs outperforming TTTC and TTTG crRNAs at later generations. On this basis, we evaluated a practical 30-60% GC range and preference for TTTA/TTTT PAMs as candidate crRNA-selection criteria within the empirically defined TSS window. Because these criteria were derived from the same screening dataset, this analysis was intended to assess their potential utility for prioritization rather than to validate the identified rules. Sequence-based filtering generally shifted the retained crRNA population toward stronger depletion, although the contributions of individual criteria were unequal (Fig. 4C, Fig. S9). Restricting crRNAs to >30% GC alone produced little improvement, consistent with the limited representation of very low-GC crRNAs in the screen. In contrast, excluding the less-preferred TTTC and TTTG PAMs produced a clearer improvement, particularly when both PAM classes were removed (Fig. S9). Combining PAM selection with exclusion of crRNAs above 60% GC produced among the largest and most consistent shifts toward stronger depletion across target genes (Fig. 4C, S9). These observations support a practical preference for intermediate GC content and TTTA/TTTT PAMs within the −260 to +50 bp targeting window.

Finally, to test if crRNA length influenced *Y. lipolytica* CRISPRi efficacy, we investigated the subset of target sites represented by complete matched N18, N20, N23 and N25 crRNAs. crRNA length had a strong effect on efficacy after accounting for target gene and matching crRNAs by target site (p = 6.2 x 10^-12^); Fig. 4D), with adjusted efficacy decreasing progressively with increasing crRNA length. Because this finding was unexpected, we considered whether it could instead arise from differences already present at T0. For example, highly active crRNAs targeting essential genes might have undergone depletion before T0 sampling, whereas less active crRNAs would remain more abundant and retain greater apparent capacity for subsequent depletion. We therefore repeated the matched crRNA-length analysis separately for essential and conditionally essential genes. The preference for shorter crRNAs was retained for both essential (*p* = 1.8 x 10^-7^) and conditionally essential (*p* = 1.6 x 10^-4^) genes (Fig. S10). Because crRNAs targeting conditionally essential genes were not expected to experience depletion before T0, the persistence of the same length-dependent trend in this subset argues against pre-T0 depletion of highly active essential-gene guides as the sole explanation. Consistent with this, T0 abundance did not differ systematically among target-matched N18, N20, N23 and N25 crRNAs (*p* = 0.71). Together, these analyses support a genuine association between shorter crRNAs length and increased CRISPRi efficacy in our dataset.

### dHyperLbCas12a outperforms both dLbCas12a and denAsCas12a in *Y. lipolytica*

The preceding analyses identified crRNA features associated with efficient CRISPRi using dLbCas12a-Mxi1, which emerged from our pooled screens as the most robust of the initially tested Cas12a variants. We next asked whether CRISPRi/a performance could be further improved with dHyperLbCas12a, an engineered LbCas12a variant containing activity-enhancing mutations that increase activity and transcriptional modulation in mammalian cells^29^ and *Candida albicans*^30^. To enable a direct comparison, we repeated the same benchmark assays used in Fig. 1. Relative to their respective non-targeting controls, strains expressing a *ZWF1*-targeting crRNA exhibited a 43% reduction in growth and 35% reduction in maximal OD in the dHyperLbCas12a-Mxi1 background, compared with reductions of only 16% and 10%, respectively, in the dLbCas12a-Mxi1 background (Fig. 5A, Table S2). The hyperactivating mutations also enhanced CRISPRa activity. dHyperLbCas12a-VPR produced a stronger *ACC1* activation phenotype, resulting in higher Nile Red fluorescence than dLbCas12a-VPR (Fig. 5B). Together, these results identify dHyperLbCas12a as the highest-performing Cas12a variant evaluated in this study and support its use for future CRISPRi/a applications in *Y. lipolytica*.

**Figure 5.**
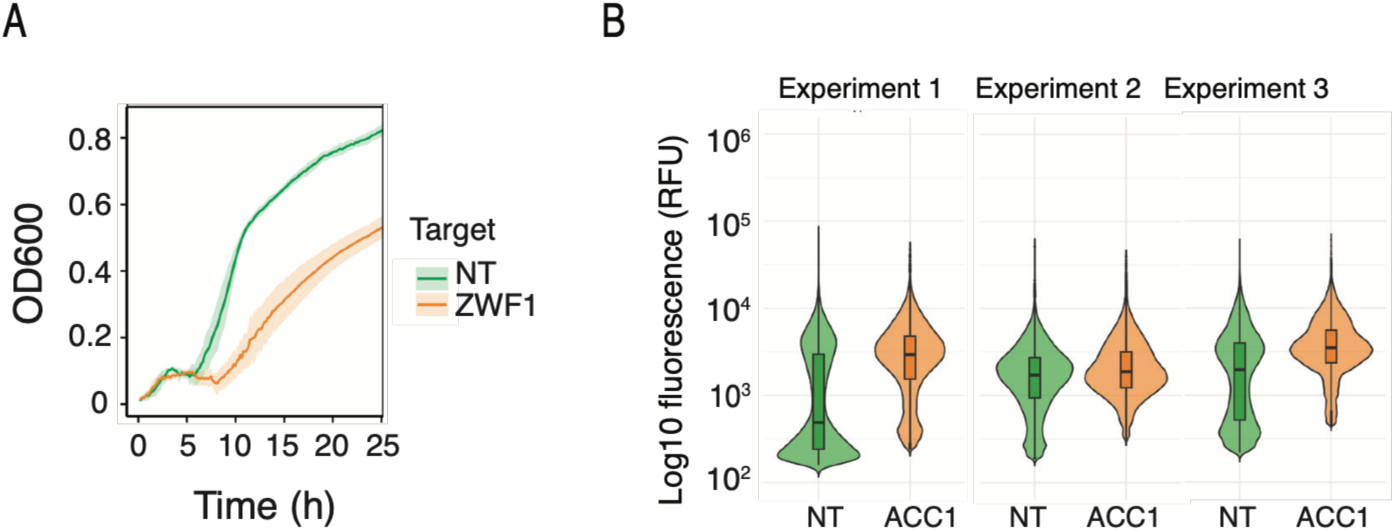
dHyperLbCas12a enables highly efficient CRISPRi/a in *Y. lipolytica*. **(A)** CRISPRi-mediated repression of *ZWF1*. Growth curves of strains expressing non-targeting (NT) or *ZWF1*-targeting crRNAs in the dHyper-Mxi1 background. Experiments were performed independently two times. Within each experiment, four biological replicates were analyzed, each consisting of three technical replicates. Technical replicates were averaged prior to analysis, and the plotted values represent the mean of the biological replicates across the two independent experiments. Shaded ribbons indicate ±1 standard deviation. **(B)** dHyperLb-based CRISPRa-mediated transcriptional activation of *ACC1*. Intracellular lipid accumulation was quantified by Nile Red staining and flow cytometry in strains expressing non-targeting (NT) or *ACC1*-targeting crRNAs. Violin plots show single-cell fluorescence distributions after exclusion of events overlapping with the corresponding unstained controls. The central line indicates the median fluorescence, and the box represents the interquartile range. Each experiment consisted of four biological replicates performed on the same day, and cells from biological replicates were pooled within each experiment for visualization. Data shown are representative of three independent experiments

## Discussion

Here, we established dCas12a-based CRISPRi and CRISPRa for pooled screens in *Y. lipolytica* and systematically compared two alternative Cas12a orthologs across both single-target and pooled assays. Initial single-gene benchmarking suggested stronger CRISPRi activity with dLbCas12a, and this trend was broadly supported in the larger pooled screen, where dLbCas12a showed more consistent performance across a broader set of targets. Both dLbCas12a and denAsCas12a nevertheless supported robust transcriptional repression, although their relative performance depended on the target, crRNA and duration of selection. dLbCas12a ultimately produced more consistent late-stage depletion across the tested genes, whereas denAsCas12a showed evidence of stronger or earlier activity for a subset of crRNAs. We subsequently evaluated the further engineered dHyperLbCas12a variant using matched CRISPRi and CRISPRa benchmark assays and found substantially stronger phenotypes than with dLbCas12a in both cases. This identifies dHyperLbCas12a as the strongest variant evaluated in this study. Its increased CRISPRi activity is consistent with the design of hyperLbCas12a, which was engineered to enhance DNA interaction^31^. Recent studies in mammalian cells^32^ and *C. albicans*^30^ have similarly identified dHyperLbCas12a as a high-performing effector for repression and activation, respectively, suggesting that its increased regulatory activity translates across divergent eukaryotic hosts.

An important finding from the pooled comparison was that differences between dLbCas12a and denAsCas12a were not explained simply by one variant being uniformly more active than the other. Both produced clear depletion phenotypes, but the identity and magnitude of the strongest crRNAs differed substantially between variants, particularly in minimal medium. This target-dependent behavior is consistent with emerging evidence that Cas12a orthologs and engineered variants can have distinct sequence preferences^27,33^. Large-scale comparisons of Cas12a nucleases in human cells have shown relatively poor correlations in activity between different orthologs at identical target sequences, despite much stronger correlations among variants derived from the same ortholog, indicating that sequence determinants of high activity are at least partly ortholog-specific^33^. Thus, a crRNA optimized for one Cas12a protein should not necessarily be expected to perform equivalently with another. This interpretation is also consistent with our findings from a smaller set of targets in *S. cerevisiae*^27^. More broadly, these results argue that evaluation of Cas12a orthologs on just a single target can be misleading. Although the stronger *ZWF1* phenotype initially observed with dLbCas12a correctly anticipated its more robust pooled performance compared to enAsCas12a in our experiments, the pooled screen revealed considerably more complex crRNA-specific differences between the two orthologs.

A limitation of the present system is that dCas12a and the crRNAs are constitutively expressed. Consequently, repression begins during library construction and expansion rather than at the nominal start of the pooled experiment. The depletion of essential-gene crRNAs already observed at G0 demonstrates that this effect is biologically relevant. This is a general concern for pooled CRISPRi screens of essential genes, because highly effective crRNAs can be selectively lost before the experimental baseline is collected. Indeed, recent fungal CRISPRi screens have introduced inducible effector expression specifically to avoid premature depletion of essential-gene perturbations^25,34^. In our experiments, both dLbCas12a and denAsCas12a nevertheless produced strong and reproducible depletion after G0, and baseline depletion was broadly comparable between orthologs, arguing against pre-depletion as the main explanation for ortholog-specific differences. For future genome-scale applications, however, inducible control of dCas12a expression would likely improve retention of strongly deleterious perturbations and expand the dynamic range available for essential-gene screening.

The deliberately permissive library design also allowed us to ask whether simple crRNA features could help prioritize more effective CRISPRi crRNAs. These analyses should be viewed as exploratory, as only a small number of target genes were examined, and efficacy was inferred indirectly from fitness-associated depletion rather than measured directly as target-gene repression. The identified features therefore primarily describe crRNAs associated with the strongest dropout in the specific context of our screen. Nevertheless, several informative patterns emerged. The crRNAs mediating strongest depletion were concentrated around and immediately upstream of the TSS, consistent with previous CRISPRi studies in microbial and mammalian cells showing preferential repression from promoter-proximal targets^13,15,23,24,35^. While several genes exhibited a dominant TSS-proximal activity peak, others showed additional regions of elevated activity within the broader promoter-proximal region, both up-and downstream of the major peak. As expected, however, position alone did not fully explain crRNA performance, and several crRNAs within otherwise favorable promoter regions showed little or no depletion. Among the optimally positioned crRNAs, intermediate GC content showed the clearest sequence-associated relationship with efficacy, whereas PAM identity and targeted strand showed weaker trends. GC content has not typically been reported as a major determinant of CRISPRi activity, but the reduction observed for high-GC crRNAs is broadly compatible with the common practice of avoiding extreme GC content in design. The preference for TTTA and TTTT over TTTC and TTTG PAMs should be interpreted cautiously, given the limited number of target genes and crRNAs. Notably, TTTT-PAM crRNAs were among the more active in our dataset, despite TTTT generally being less preferred by LbCas12a^36,37^. Interestingly, large-scale Cas12a cleavage screens in human cells indicate slightly lower activity at TTTA than at TTTC or TTTG PAMs^33^, whereas TTTA was among the more active PAMs in our CRISPRi dataset. This difference may reflect distinct PAM preferences for transcriptional repression versus nuclease-mediated cleavage, or differences between host systems. More broadly, our observation that activity varies among individual PAM sequences is consistent with large-scale Cas12a datasets showing PAM-dependent differences even within the canonical TTTV class^27,33^.

Sequence context beyond the canonical PAM can also influence Cas12a activity. Previous cleavage and precision editing studies have identified effects of the nucleotide immediately upstream of the PAM^27,33^, and of potential base-pairing interactions between the DR and the variable part of the crRNA^38^. We did not detect an association between either of these features and CRISPRi efficacy in our dataset, suggesting that any such effect is either small in this context or not well resolved by the available crRNAs. Finally, the lack of predictive value of CHOPCHOP rank and efficiency in our dataset illustrates the limited transferability of general design scores to a specific CRISPRi system. CHOPCHOP v3 incorporates Cas12a efficiency scoring and provides targeting modes for transcriptional regulation^28^, but these scores were not developed specifically for dLbCas12a-Mxi1-mediated repression in *Y. lipolytica.* Together, the GC- and PAM-associated trends observed here are therefore best viewed as empirical criteria that may help prioritize crRNAs for this system, rather than as universal design rules.

Spacer length produced one of the strongest and most unexpected associations in the screen, with N18 crRNAs consistently outperforming matched N20, N23 and N25 variants. Because spacer variants targeted exactly the same genomic positions, this comparison is less susceptible to confounding by position than the other sequence-feature analyses. The persistence of the effect in both essential and conditionally essential genes, together with the absence of a systematic spacer-length difference in G0 abundance, also argues against pre-screen depletion as its sole explanation. Shorter Cas12a spacers are mechanistically plausible: early biochemical work showed that approximately 18 nucleotides of complementarity are sufficient for efficient Cas12a cleavage^36^, and efficient dCas12a-mediated repression in bacteria can also be maintained with shortened guide sequences^39^. However, spacer-length effects are not universal. Experiments with FnCas12a have shown clear target dependence, with 18-nt guides outperforming longer guides at some loci but not others^40^. This target and effector dependence is also consistent with our own Cas12a cleavage experiments in *S. cerevisiae*, where 18-nt spacers supported efficient editing with enAsCas12a but were less effective with FnCas12a ^27^. The consistency of the effect across matched target sites is nevertheless striking and supports further validation using an independent set of genes and crRNAs.

Because our pilot library contained substantially more CRISPRi than CRISPRa targets, the CRISPRa dataset was considerably smaller and therefore did not permit a similarly comprehensive analysis of crRNA features. Nevertheless, the enrichment patterns observed for *ERG11* were broadly consistent with the promoter-proximal targeting preference identified for CRISPRi. At the same time, enriched *ERG11*-targeting crRNAs were distributed across a broader promoter region than the strongly depleted crRNAs observed in the CRISPRi fitness screens. This likely reflects the nature of positive selection rather than a fundamentally broader targeting window: under stringent fluconazole selection, even a modest increase in *ERG11* expression could provide a substantial relative growth advantage when most other cells are not growing. Consequently, crRNAs producing relatively small transcriptional changes can become strongly enriched over multiple generations. This may explain why CRISPRa appeared active across a broader positional range and suggests that the effective targeting window can depend on the sensitivity of the phenotypic readout.

Together, these results establish Cas12a as a versatile platform for both CRISPRi and CRISPRa screening in *Y. lipolytica* while also illustrating why effector and crRNA design need to be optimized in the relevant host and assay context. dLbCas12a provided robust pooled performance despite constitutive expression, and empirical analysis identified a promoter-proximal targeting interval together with sequence and spacer-length features that can guide future crRNA prioritization. The substantially stronger phenotypes obtained with dHyperLbCas12a further suggest that effector engineering can provide an additional, and potentially larger, gain in regulatory activity than crRNA optimization alone. Because dHyperLbCas12a is derived from LbCas12a, the crRNA preferences identified here for dLbCas12a may also provide a useful starting point for guide design with the hyperactive variant. This possibility is supported by high-throughput nuclease profiling showing that target-specific activity is considerably more correlated among engineered variants derived from the same Cas12a ortholog than between distinct Cas12a orthologs^33^. However, this remains to be tested directly, because the hyperactivating mutations could alter the relative performance of individual crRNAs. Future work combining dHyperLbCas12a with inducible expression and independently validated crRNA-design criteria should provide a strong foundation for larger-scale functional and metabolic screens in *Y. lipolytica*. More broadly, the combination of an improved effector with empirically informed library design should increase the fraction of functional perturbations per library and thereby reduce the library size required for robust pooled screening. Together, these advances substantially expand the functional-genomics toolbox available for this industrially relevant yeast and provide a basis for future genome-scale gain-and loss-of-function studies.

## Materials and methods

### Media, strains, plasmids and transformations

Plasmids and strains used in this study are listed in Table S3 and Table S4, respectively. DH5α *Escherichia coli* was used for plasmid construction and propagation and was cultured in Lysogeny broth (LB) supplemented with 100 mg/L ampicillin in broth or plates. Plasmid library-construction used E. cloni® 10G Supreme Electrocompetent Cells (Biosearch Technologies, Petaluma, CA, USA) according to the manufacturer’s instructions. Yeast strains were cultivated at 30°C in YPD (10 g L⁻¹ yeast extract, 20 g L⁻¹ peptone, and 20 g L⁻¹ glucose), SC (6.7 g L⁻¹ yeast nitrogen base without amino acids supplemented; 2.002 g L⁻¹ SC Complete Drop-Out mixture [Formedium DSCK]; 20 g L⁻¹ glucose), SC-Arg-Lys (SCAL, 6.7 g L⁻¹ yeast nitrogen base with ammonium sulfate and without amino acids ; with 6.7 g L⁻¹ Drop-Out Mixture Minus Arginine and Without Lysine [Formedium FMA23]; 20 g L⁻¹ glucose), SCM (6.7 g L⁻¹ yeast nitrogen base with ammonium sulfate and without amino acids supplemented; 20 g L⁻¹ glucose), SCGly (6.7 g L⁻¹ yeast nitrogen base without amino acids supplemented; 2.002 g L⁻¹ SC Complete Drop-Out mixture [Formedium DSCK]; 50 g L⁻¹ glycerol), SCXyl (6.7 g L⁻¹ yeast nitrogen base without amino acids supplemented; 2.002 g L⁻¹ SC Complete Drop-Out mixture [Formedium DSCK]; 0.25 g L⁻¹ glucose; 5.85 g L⁻¹ xylose) or fermentation medium (1.9 g L⁻¹ yeast nitrogen base without ammonium sulfate and amino acids, 0.61 g L⁻¹ urea, 60 g L⁻¹ glucose, and 0.01 g L⁻¹ iron sulfate; initial pH 6.5). Unless specified otherwise, yeast transformations were performed using a lithium acetate/PEG protocol as described in Qi *et al*. ^41^

### Plasmid design, cloning and CRISPRi/a strain construction

CRISPRa and CRISPRi systems were based on a two-plasmid architecture comprising an integrative dCas12a effector cassette and a replicative crRNA expression plasmid (Fig. 1A). Three nuclease-deficient Cas12a variants were evaluated: denAsCas12a (E993A), dLbCas12a (D832A), and dLbHyperCas12a, which additionally carries the activity-enhancing mutations D156R, D235R, E292R, and D350R^29^. All coding sequences were codon-optimized for *Y. lipolytica.* dCas12a expression cassettes were assembled by placing each dCas12a variant under the control of the constitutive promoter pUAS1B8-TEF(136) and the CYC1 terminator in plasmid pCbf6578. Transcriptional effector domains were subsequently fused to the C-terminus of dCas12a to generate CRISPRa (VPR; amplified from pCRISPRa_VPR_yl, Addgene #107677, from Schwartz *et al*.^8^) and CRISPRi (Mxi1; amplified from pCRISPRi_Mxi1_yl, Addgene #91248, from Schwartz *et al*.^9^) constructs.

crRNA expression vectors were derived from pCfB3405 (Addgene #106166, from Holkenbrink *et al*.^21^) by replacing the *POT1* promoter with a synthetic crRNA expression cassette comprising the SCR1’-tRNAGly promoter, the corresponding Lb/AsCas12a direct repeat sequence, a SapI restriction site, a 23-nt filler sequence containing an XhoI restriction site, a second SapI restriction site, and a poly(T) terminator. Individual crRNAs were ordered as complementary DNA oligonucleotides (forward and reverse strands), annealed to generate double-stranded inserts, and cloned into BspQI-digested crRNA expression vectors by Golden Gate Assembly. Sequences of crRNAs targeting *ZWF1* and *ACC1* in experiments described in Fig.1 and Fig.5 are listed in Table S5. All plasmids are available upon request.

dCas12a expression cassettes were integrated into the neutral genomic locus IntC_2^21^ using hygromycin selection. Correct integration was verified by colony PCR. The hph marker, flanked by loxP sites, was subsequently excised by transient expression of Cre recombinase, and marker loss was confirmed by PCR and hygromycin sensitivity.

### Growth curves

Growth assays were performed in 96-well plates using a Synergy H1 multimode reader (BioTek). Strains were first cultivated overnight in 5 mL SC medium supplemented with 250 μg mL⁻¹ clonNAT in glass culture tubes mounted on a rotating wheel at 30°C. Cells were subsequently harvested by centrifugation, washed once with sterile water, and used to inoculate fresh medium in Nunclon 96-well round-bottom plates to an initial OD_600_ of 0.2. Plates were incubated at 30°C with continuous linear shaking and growth was monitored by measuring OD_600_ every 10 min for 24 h. OD600 values were subsequently used to calculate growth parameters including maximum OD_600_ (maxOD) and area under the growth curve (AUC) using a custom R script.

### Flow cytometry

For lipid quantification in individual strains, cells were grown overnight in 5 mL SC medium supplemented with 250 μg mL⁻¹ clonNAT in glass culture tubes and incubated on a rotating wheel at 30°C. After approximately 16-18 h of cultivation, cells were pelleted and stained with Nile Red. Briefly, 10⁶-10⁷ cells were collected by centrifugation, washed once with PBS, and incubated for 15 min in the dark with 4 μg mL⁻¹ Nile Red prepared from a 1 mg mL⁻¹ stock solution in acetone. Excess dye was removed by two washes with PBS, after which cells were resuspended in focusing fluid and analyzed immediately using an Attune NxT flow cytometer (Thermo Fisher Scientific). Nile Red fluorescence was measured in the YL1-A channel, and 200,000 events were recorded per sample. An unstained control was included for each strain and condition to define background fluorescence. Events overlapping with the corresponding unstained control population were excluded from subsequent analyses. Fluorescence distributions were visualized as violin plots generated using a custom R script. Each experiment consisted of three independent biological replicates, each containing four technical replicates.

### Design of crRNA library

To maximize the diversity of perturbations evaluated, 13 target genes were selected across a wide range of biological functions and selection phenotypes, including drug resistance, condition-dependent growth, and lipid accumulation. For each target gene, crRNAs were designed on the *Y. lipolytica* W29 reference genome and annotation available from Ensembl Fungi (assembly GCA_001761485). Target regions extended from 1 kb upstream of the start codon up to the first 10% of the total gene length, considering crRNAs on both the template and non-template strands. Candidate crRNAs were required to have a TTTV PAM, a GC content of 20 -85%, lack homopolymeric stretches exceeding five consecutive As or Ts and 3′-proximal T4 stretches to prevent premature PolIII transcription termination, and lack BspQI/SapI restriction sites. CrRNAs were further filtered based on predicted off-target potential using Cas-OFFinder^42^. Only crRNAs whose closest predicted off-target site contained at least four mismatches for TTTV PAMs or three mismatches for TTV PAMs were retained. In addition, the five highest-ranked CHOPCHOPv3^28^-designed crRNAs per gene were included when not already represented in the library, which led to inclusion of some TTTT PAM crRNAs. To assess the impact of spacer length on targeting efficacy, crRNAs targeting these highest-ranked sites were designed with spacer lengths of 18, 20, 23, or 25 nt. 32 non-targeting (NT) crRNAs of 23 nt length were included as negative controls. NT candidate sequences were generated by random sampling of all four nucleotides (A, C G, T), while restricting homopolymer stretches to a maximum of five consecutive identical nucleotides. To ensure sequence diversity, a minimum Levenshtein distance of nine was required between any two NT crRNAs. NT crRNA candidate sequences were subsequently screened against the W29 reference genome using BLAST+ to exclude potential genomic targets, retaining only sequences whose closest genomic match differed by at least eight mismatches or insertions/deletions. From the remaining candidates, 32 non-targeting crRNAs were randomly selected for inclusion in the library. The final pilot library comprised 752 crRNAs, including the 32 non-targeting controls. For oligonucleotide library synthesis, crRNAs were arranged in 200-nt oligonucleotides, each containing four individual crRNA sequences separated by BspQI sites, allowing individual crRNAs to be released by BspQI digestion. Oligonucleotide pools were synthesized by Twist Bioscience.

### Cloning and transformation of the pooled library

The pooled oligonucleotide library was amplified using KAPA HiFi HotStart ReadyMix (Kapa Biosystems) with 0.5 ng of template DNA and a maximum of 14 PCR cycles to limit amplification bias. Four independent 50 µL PCR reactions were performed in parallel and pooled following amplification to minimize PCR jackpotting. PCR products were purified using AMPure magnetic beads (MAGBIO Genomics) at a 1.8X bead-to-sample ratio. The amplified oligonucleotide pool was digested with BspQI to release individual crRNA inserts. Briefly, 1 μg of PCR-amplified oligonucleotide pool was digested overnight at 50°C in CutSmart buffer, followed by dephosphorylation using QuickCIP phosphatase (NEB). After heat inactivation, the released crRNA inserts were used directly for downstream cloning without further purification. The crRNA expression plasmid backbone was prepared by sequential digestion with BspQI and XhoI. BspQI digestion linearized the plasmid and excised the filler sequence located between the two cloning sites. Subsequently, XhoI was used to digest the released filler fragment, thereby reducing background colonies arising from vector self-ligation during library cloning. The linearized backbone was purified by agarose gel electrophoresis extraction. Digested crRNA inserts (10 ng) and plasmid backbone (500 ng) were ligated using T4 DNA ligase, with a 1h incubation at 25°C followed by enzyme inactivation at 65°C for 10 min. Ligation products were subsequently desalted by drop dialysis^43^ prior to electroporation into E. cloni 10G Supreme electrocompetent *E. coli* (BioSearch Technologies). Following 1 h recovery at 37°C, transformed cells were plated on LB agar supplemented with 100 μg mL⁻¹ carbenicillin. Transformation efficacy and library coverage were estimated from dilution plates, and libraries were recovered only when the number of transformants exceeded a minimum coverage of 100X per crRNA. Bacterial colonies were harvested by plate washing with LB supplemented with carbenicillin and thoroughly homogenized to ensure even representation of clones. Aliquots were preserved as glycerol stocks at -80°C at a final OD_600_ of 30, while the remaining cells were used for plasmid extraction using a modified MiraPrep protocol. Purified plasmid libraries were subsequently stored at −20°C until use. To assess library integrity, individual colonies were screened by colony PCR and agarose gel electrophoresis to distinguish correctly assembled plasmids from empty-vector background. Selected positive clones were further validated by Sanger sequencing to confirm the presence and identity of the inserted crRNA sequences.

Yeast library transformations were performed according to Robertson *et al.*^44^ with minor modifications. Briefly, following heat shock, cells were collected by centrifugation and resuspended in 1 mL YPD. Transformants were allowed to recover for 4 h at 30°C in a thermomixer shaking block (500 rpm). Following recovery, all transformation reactions were pooled, pelleted, and resuspended in 1 mL YPD. To estimate transformation efficacy, serial dilutions (0.1%, 0.01%, and 0.001%) were plated onto YPD supplemented with 250 µg/mL clonNAT. The remaining cells were spread onto large square YPD-clonNAT plates to maintain library coverage. After 2 days of incubation at 30°C, colonies were harvested by washing the plates with YPD, thoroughly homogenized, and stored as multiple glycerol stocks at −80°C at a final OD_600_ of 15.

### Screening of the CRISPRi and CRISPRa libraries

Throughout all screening experiments, cultures were maintained at a minimum coverage of 1,000X and kept in exponential growth phase to ensure continuous CRISPR activity and preserve library representation. For each screening experiment, a preculture step was performed by growing yeast libraries from glycerol stocks in YPD supplemented with 250 μg mL⁻¹ clonNAT for approximately five generations. Following recovery, four independent cultures were inoculated in the corresponding screening medium, each supplemented with 250 μg mL⁻¹ clonNAT to maintain selection of the crRNA plasmid. Cultures were propagated for 25 generations, with samples collected every five generations (G5, G10, G15, G20, and G25). At each time point, cells were harvested by centrifugation and the remaining culture was back-diluted into fresh medium to maintain exponential growth. dLbCas12a-CRISPRi libraries were screened in SCC, SCM, SC-Arg-Lys, and SC-Arg-Lys supplemented with 0.156 μg mL⁻¹ L-canavanine. dLbCas12a-CRISPRa libraries were screened in SCC and SCC supplemented with 2 μg mL⁻¹ fluconazole. denAsCas12a-CRISPRi libraries were screened in SCC and SCM. For dLbCas12a-based CRISPRa screens of lipid accumulation phenotypes, library glycerol stocks were incubated in fermentation medium for 5 days at 30°C. Following cultivation, cells were stained with Nile Red, as described above, and analyzed using a BD FACS Aria Fusion cell sorter. Events were first gated based on forward scatter area (FSC-A) and side scatter area (SSC-A) to exclude debris and non-cellular particles. Doublets were subsequently removed using FSC-H versus FSC-A and SSC-H versus SSC-A gating, retaining only singlet events. A final FSC/SSC gate was applied to select a homogeneous population of intact yeast cells for fluorescence analysis. Intracellular lipid content was quantified using Nile Red fluorescence (YL1-A channel). The final singlet population was partitioned into four gates spanning increasing Nile Red fluorescence intensities, corresponding to progressively higher intracellular lipid content. Cells from each fluorescence gate were sorted independently and collected for downstream plasmid extraction and crRNA sequencing. For subsequent analyses, crRNA enrichment was quantified by comparing crRNA abundances in the higher-fluorescence gates (G2-G4) relative to the lowest-fluorescence gate (G1).

### Yeast plasmid extraction, crRNA amplification and NGS

Plasmid DNA was extracted from pooled *Y. lipolytica* cultures using the QIAprep Spin Miniprep Kit according to the manufacturer’s protocol for yeast, with several modifications. Briefly, cell pellets were resuspended in 600 μL of 1 M sorbitol, 0.1 M EDTA (pH 7.4) containing 200 U zymolyase-100T and incubated for 30 min at 30°C to promote cell wall digestion. Following addition of 600 μL Buffer P1 supplemented with RNase A (0.1 mg mL⁻¹), cells were mechanically lysed using acid-washed glass beads (425-600 μm) in a FastPrep instrument (2 × 30 s at 6 m s⁻¹, with 1 min on ice between cycles). After removal of the beads, 1.2 mL Buffer P2 and 1.2 mL Buffer N3 were sequentially added. Lysates were subsequently clarified by centrifugation and processed using QIAprep Spin Miniprep columns according to the manufacturer’s protocol.

crRNA sequences were amplified from plasmid DNA using a two-step PCR strategy. In the first PCR, the crRNA-containing region was amplified using the F/R (5’◊3’) primers CTTTCCCTACACGACGCTCTTCCGATCTCCGGTTCGATTCCGGGTC/ GGAGTTCAGACGTGTGCTCTTCCGATCTGATCGCACGCATTCCGTTG. Reactions were performed in four parallel reactions containing RepliQa polymerase master mix, 5 ng template DNA, and 5 μM primers. Amplification was carried out for 24 cycles, and PCR products were pooled and purified by two-sided AMPure bead size selection (350-450 bp). Illumina adapters and dual indexes were added in a second PCR. Four parallel 30 μL reactions containing up to 10 ng purified PCR1 product were amplified for 12 cycles, yielding a final amplicon of approximately 440 bp. Indexed libraries were purified by double-sided AMPure bead size selection (350-450 bp), quantified using the Qubit dsDNA High Sensitivity Assay (Thermo Fisher Scientific), and pooled equimolarly. Sequencing was performed on an AVITI system (Element Biosciences) using paired-end 150 bp reads (2 × 150 bp).

### Processing of crRNA sequencing data

crRNAs were extracted from reads using a custom R script using fuzzy matching to constant flanking sequences to identify start and end of each crRNA, then tallied per sample and matched to the designs based on sequence. For growth-based screens, the R package DESeq2^45^ was used for normalization and statistical inference of changes in crRNA abundance over time. Size factors were estimated from NT crRNAs after excluding sparsely represented controls with more than five zero-count observations, leaving 27–31 NT crRNAs for normalization. Abundance changes were estimated separately within each Cas ortholog and condition. Sampling generation was treated as a factor with generation 0 (G0) as the reference, and biological replicate was included as a blocking factor. crRNAs with fewer than five reads across the dataset were removed before dispersion estimation and model fitting. For each post-G0 generation, a Wald contrast against G0 was extracted, and log_2_ fold changes (log_2_FCs) were additionally shrunk using lfcShrink(…, type = “ashr”); the shrunk log_2_FC was used as the primary reported effect size. For analyses aimed at defining the preferred TSS-targeting window and identifying crRNA features associated with CRISPRi efficacy, the same DESeq2 workflow was repeated using only dLbCas12a-Mxi1 samples from SCC and SCM. Shrunken log_2_FCs from these models were used for the positional and guide-feature analyses described below. For the FACS-based CRISPRa screen, crRNA counts were normalized to counts per million (CPM) within each sample. crRNA enrichment between fluorescence-sorted populations was then calculated as the log_2_FC in CPM between each higher-lipid gate and the corresponding baseline gate (G1).

### Analysis of features associated with CRISPRi efficacy

To identify the optimal targeting window relative to the TSS, TSSs were taken from an in-house, unpublished STRIPE-seq dataset (manuscript in preparation) generated for the W29 strain used as background for CRISPRi/a experiments. Briefly, cultures were grown in SC medium at 30°C in shake flasks, and cells were harvested during exponential growth at OD_600_ values of 0.3 and 0.8. Three biological replicates were collected for each condition and sampling point. Total RNA was isolated using the RNeasy Mini Kit (Qiagen) according to the manufacturer’s instructions. Library preparation, read processing and TSS identification followed the published STRIPE-seq protocol^46^ with one modification to enable generating dual-indexed libraries. Sequencing was performed on an AVITI platform (Element Biosciences) using paired-end 150 bp reads (2 × 150 bp). We retained TSS with a total count of at least 20 across replicates and of at least 3 in at least two replicates. We further required TSS to be located on the same strand within 2,000 bp upstream to 150 bp downstream of the start codon, or in the gene’s first intron. A gene’s dominant TSS was defined as the highest-count TSS. A TSS ≥25 bp away with ≥10% of the dominant TSS’s count and support in at least two replicates was called an alternative TSS.

The analysis of CRISPRi crRNA efficacy as a function of distance from the dominant TSS included crRNAs targeting essential and conditionally essential genes and used data from SCM at G20 relative to G0. Efficacy was defined as the negative shrunken log_2_fold change (−log_2_FC), such that stronger depletion corresponded to higher efficacy values. Only 23-nt crRNAs were included to avoid counting different crRNA lengths targeting the same genomic position multiple times. To account for differences in overall depletion magnitude between target genes, crRNA efficacy was centered within each gene by subtracting the median efficacy of all crRNAs targeting that gene. A sliding-window analysis was then performed using 100-bp windows with 25-bp increments. As library design was deliberately permissive to enable the subsequent identification of parameters influencing crRNA efficacy, we focused on relatively effective crRNAs rather than average crRNA performance in each window. Accordingly, the 80th percentile of gene-centered crRNA efficacy was calculated for each gene and window, and the median of these gene-specific values was then calculated for each window to prevent genes with more crRNAs from disproportionately influencing the positional profile. crRNA feature analysis used the same dLbCas12a-Mxi1 SCM dataset as the TSS-window analysis, restricted to 23-nt crRNAs within the empirically defined −260 to +50 bp window relative to the dominant TSS. Efficacy was again defined as −log_2_FC. A linear model was fitted including target gene, TSS distance, GC-content class, PAM sequence, targeted strand, the nucleotide immediately 5′ of the PAM, and predicted perturbation of the DR structure determined using the Vienna RNA package^47^. Target gene was included as a fixed effect, and TSS distance was modeled using a natural cubic spline (df = 3), allowing for residual positional effects within the optimal window. GC content was grouped into <30%, 30–40%, 40–50%, 50–60% and >60% in the primary analysis. As a sensitivity analysis, GC class was replaced by continuous GC content modeled using a natural cubic spline (df = 3). Feature effects were evaluated using partial F-tests (drop1(…, test = “F”)), with Holm correction applied across the tested crRNA features. Estimated marginal means and 95% confidence intervals were obtained using emmeans, with Tukey adjustment for pairwise comparisons. The same primary model was fitted independently at G5, G10, G15, G20 and G25 using crRNAs represented at all five generations. To assess the effect of crRNA length only target sites represented by complete matched sets of 18, 20, 23 and 25 nt crRNAs were included. Length was treated as a categorical variable in a linear mixed-effects model containing target gene as a fixed effect and target site as a random intercept. The overall spacer-length effect was tested using a Type II F-test with Kenward–Roger degrees of freedom, and estimated marginal means and Tukey-adjusted pairwise comparisons were obtained using function *emmeans*. To assess whether crRNA-selection metrics generated by CHOPCHOP v3 were informative for CRISPRi activity in *Y. lipolytica*, experimentally measured efficacy was compared with CHOPCHOP Rank and Efficacy scores. Rank and Efficacy were highly correlated and hence tested separately using linear models containing target gene as a fixed effect and the respective CHOPCHOP metric as a continuous predictor. The contribution of each metric was evaluated using an F-test comparing the full model with the corresponding reduced model.

## Acknowledgements

We are grateful to the VIB Nucleomics Facility for assistance with sequencing and to the VIB Flow Core Leuven for support with FACS. This work was supported by the VIB Grand Challenges program (BIOPET project).

---

**Supplementary Figure 1.**
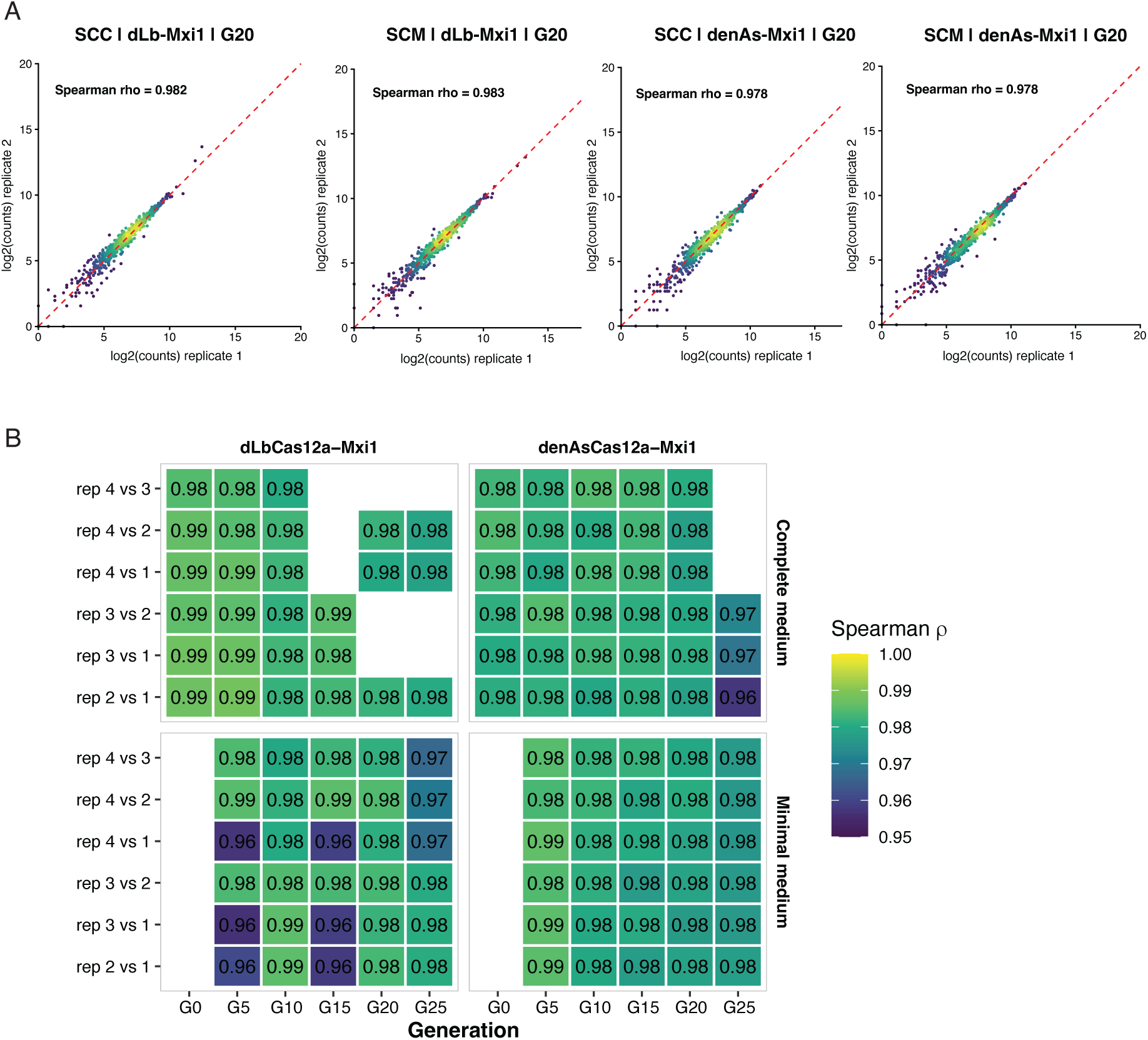
Reproducibility of pooled CRISPRi screens across biological replicates. **(A)** Representative replicate correlations at G20 for dLbCas12a-Mxi1 and denAsCas12a-Mxi1 in complete (SCC) and minimal (SCM) medium. Normalized crRNA abundances are shown as log_2_-transformed counts for biological replicate pairs; dashed lines indicate (x=y). Spearman correlation coefficients are shown within each panel. **(B)** Spearman correlations between all biological replicate pairs. Values and shading within tiles indicate Spearman ρ.

**Supplementary Figure 2.**
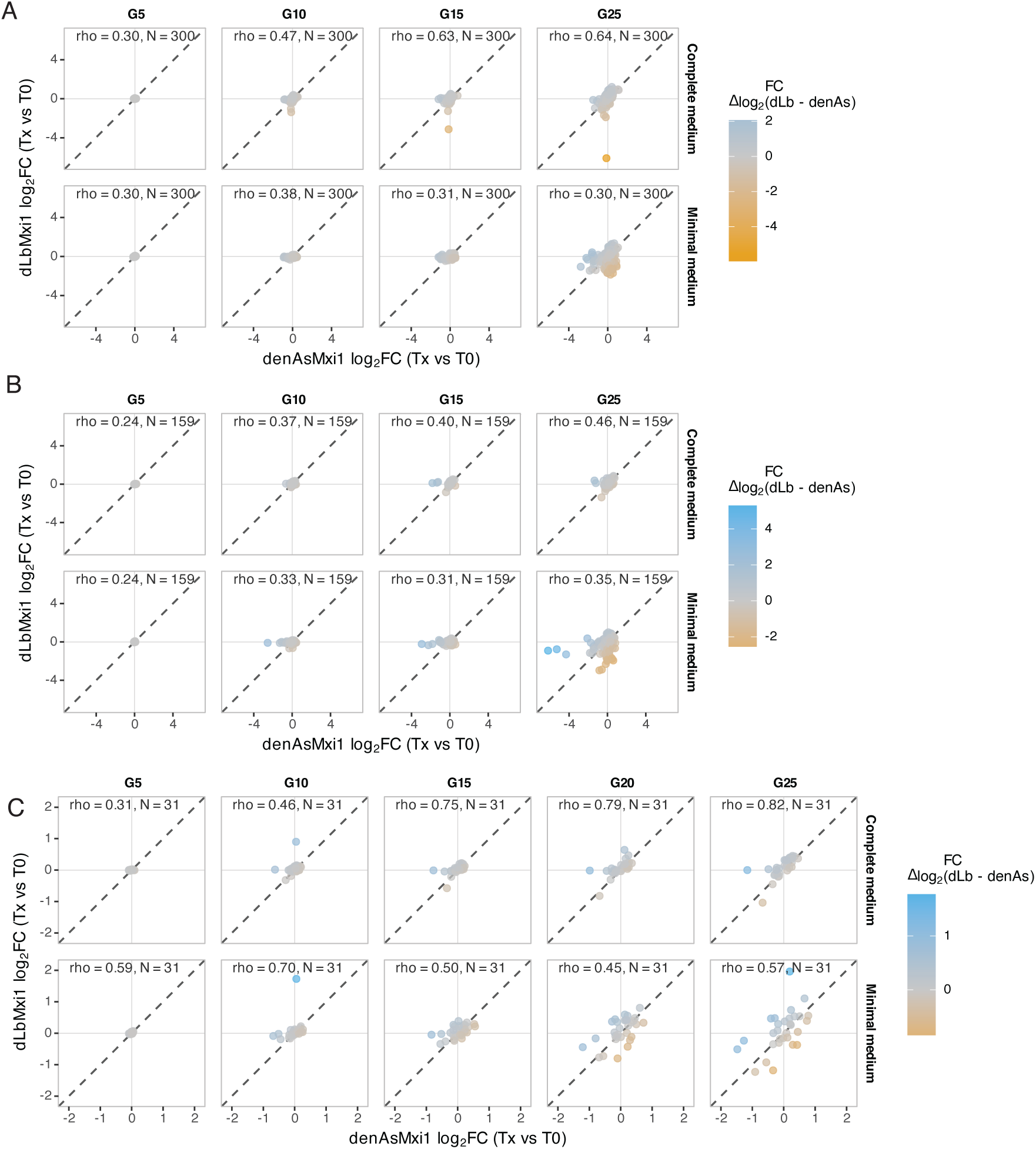
Concordance between dLbCas12a-Mxi1 and denAsCas12a-Mxi1 across generations. Matched crRNA-level log_2_-fold-changes (log_2_FCs) between denAsCas12a-Mxi1 and dLbCas12a-Mxi1 across generations. **(A)** Essential-gene crRNAs in complete and minimal medium. **(B)** Conditionally essential-gene crRNAs in complete and minimal medium. **(C)** Non-targeting crRNAs. Each point represents the same crRNA tested with both Cas variants; dashed diagonals indicate equal effects. Points are coloured by the difference in log_2_FC between Cas variants. Spearman ρ and the number of matched crRNAs are indicated in each panel. The tested timpeoint is indicated at the top of each panel.

**Supplementary Figure 3.**
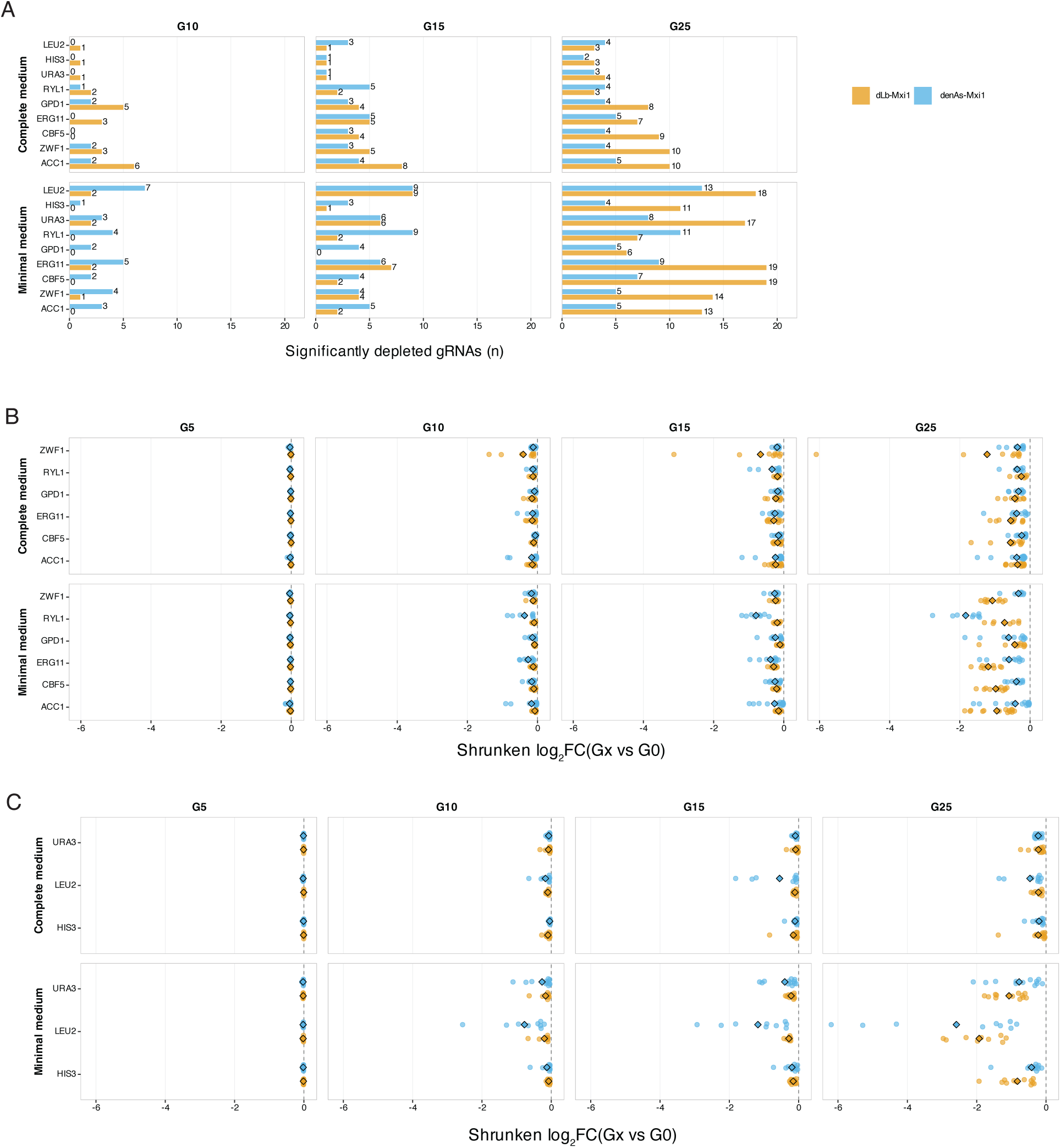
Temporal comparison of CRISPRi efficacy between Cas12a orthologs. **(A)** Number of significantly depleted crRNAs per gene for dLbCas12a-Mxi1 and denAsCas12a-Mxi1 across generations in complete and minimal medium. G5 is omitted because there were no significant crRNAs. **(B–C)** Log_2_FCs for the most depleted 20% of crRNAs within each gene, condition and Cas ortholog. **(B)** Essential genes. **(C)** Conditionally essential genes. Points represent individual crRNAs and diamonds indicate the corresponding mean log_2_FC per gene. More negative log_2_FC values indicate stronger fitness defects.

**Supplementary Figure 4.**
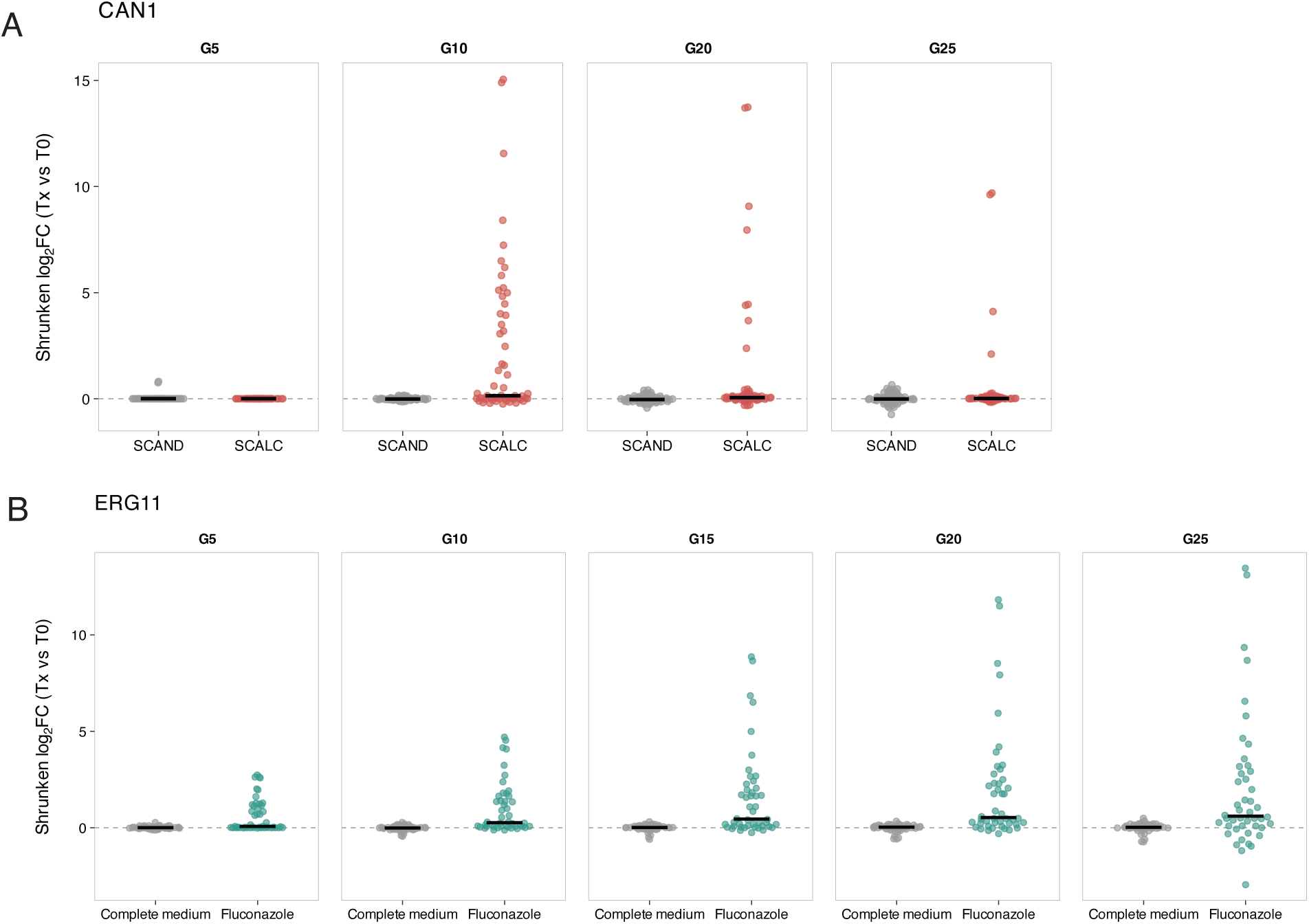
Time-resolved enrichment of drug-resistance phenotypes in CRISPRi and CRISPRa screens. **(A)** CRISPRi targeting of *CAN1* during growth in complete medium without (SCAND) or with (SCALC) L-canavanine. Each dot represents one crRNA. Enrichment of *CAN1*-targeting crRNAs under L-canavanine selection is consistent with reduced *CAN1* expression conferring resistance. **(B)** CRISPRa targeting of *ERG11* during growth in complete medium with or without fluconazole. Each dot represents one crRNA. Enrichment of *ERG11*-targeting crRNAs under fluconazole selection is consistent with increased *ERG11* expression promoting resistance. Black bars indicate medians.

**Supplementary Figure 5.**
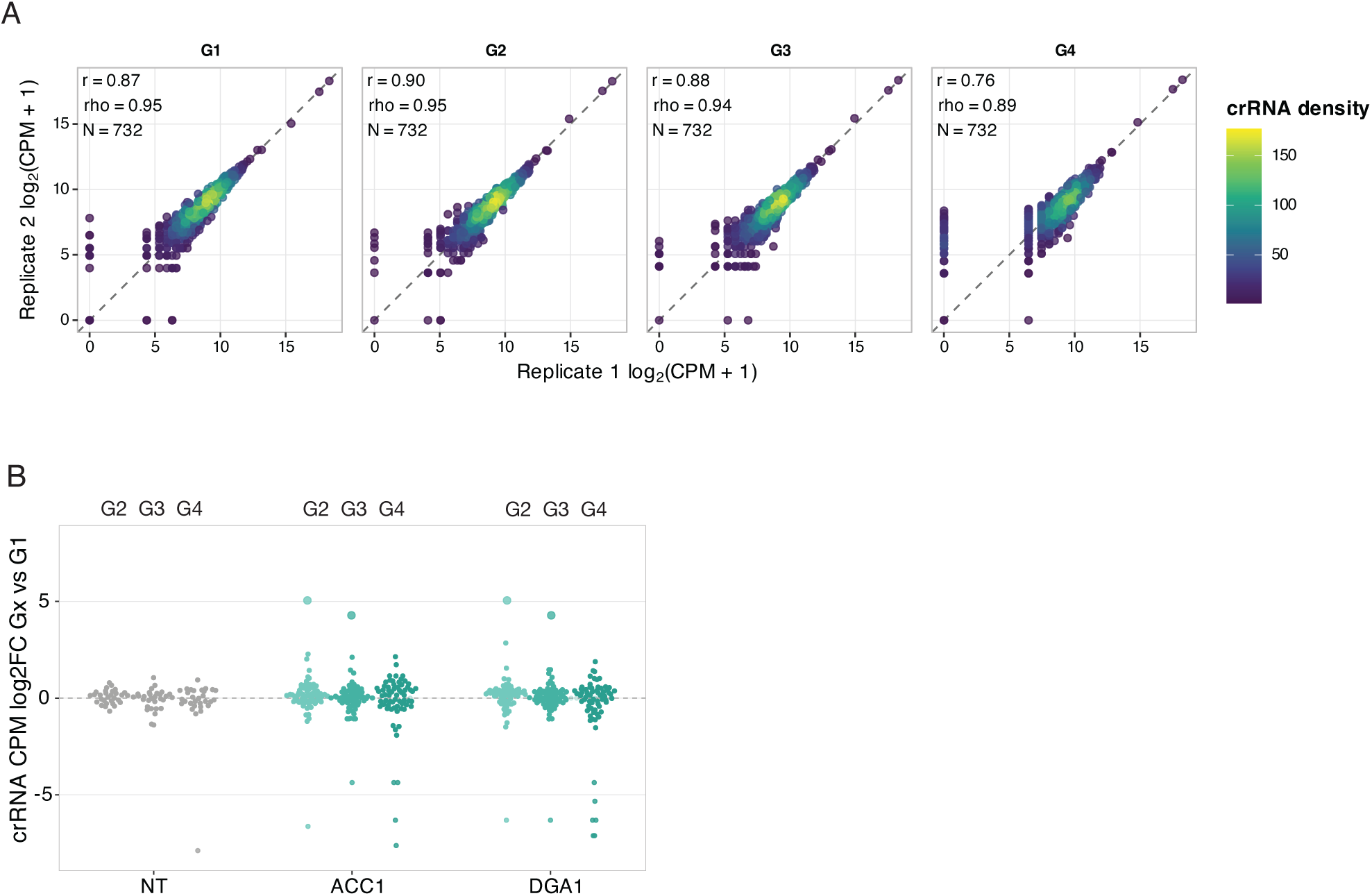
Reproducibility and crRNA enrichment in the CRISPRa lipid FACS screen. **(A)** Correlation of crRNA abundances between biological replicates for the G1, G2, G3 and G4 FACS populations. CrRNA abundance is shown as log_2_(CPM + 1); dashed lines indicate (x=y). Pearson (r), Spearman ρ and the number of crRNAs are shown in each panel. (**B)** CRISPRa screen for altered lipid accumulation following targeting of *ACC1* and *DGA1*. Cells were stained with Nile Red and sorted by FACS into four gates spanning baseline to increased lipid-associated fluorescence (G1 < G2 < G3 < G4). CrRNA enrichment relative to the baseline G1 population is shown for replicate 1 for G2-G4 (left to right) for nontargeting (grey), and *ACC1*-/*DGA1*-targeting crRNAs (in turquoise shades). Positive log2FC values indicate enrichment relative to G1. Larger points indicate crRNAs with zero counts in the G1 reference gate and non-zero counts in the corresponding higher-lipid gate.

**Supplementary Figure 6.**
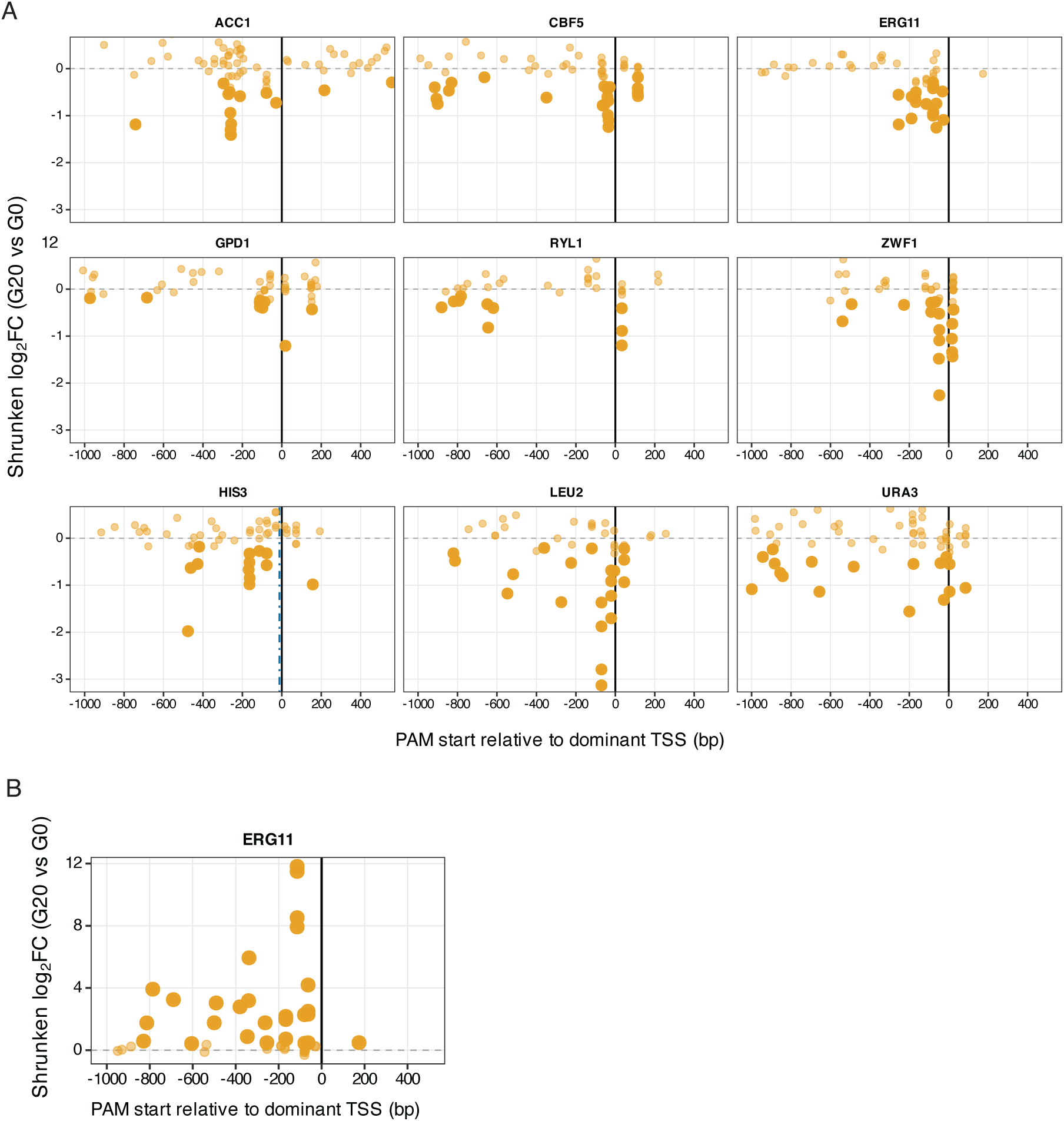
Relationship between crRNA position relative to the TSS and CRISPRi activity across target genes. crRNA log_2_ fold-changes (log_2_FCs) at G20 relative to G0 are shown as a function of distance from the dominant transcription start site (TSS) for individual essential genes in minimal medium. Each point represents an individual crRNA. Larger points indicate crRNAs with an adjusted *p* value ≤ 0.05.

**Supplementary Figure 7.**
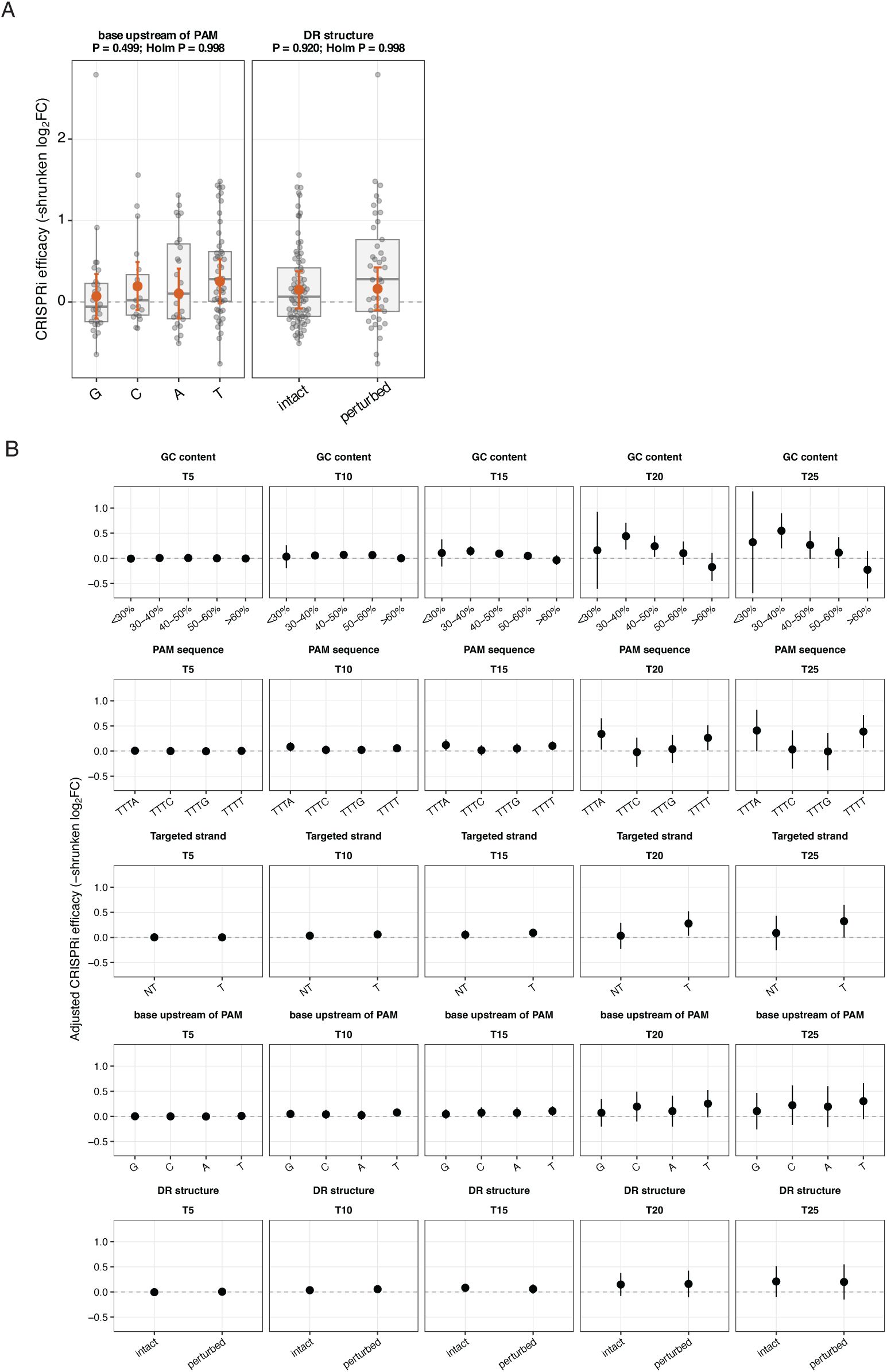
Associations between crRNA features and CRISPRi efficacy. **(A)** Associations between CRISPRi efficacy and the nucleotide immediately 5′ of the PAM and predicted perturbation of the direct-repeat (DR) structure for crRNAs within the −260 to +50 bp TSS-targeting window. Individual crRNAs are shown as grey points and boxplots; orange points and error bars indicate adjusted estimated marginal means and 95% confidence intervals from the linear model accounting for target gene, residual TSS-position effects and the other crRNA features. **(B)** Adjusted estimated marginal means and 95% confidence intervals for crRNA features across generations. The same model was fitted independently at each generation using crRNAs represented across all analyzed generations.

**Supplementary Figure 8.**
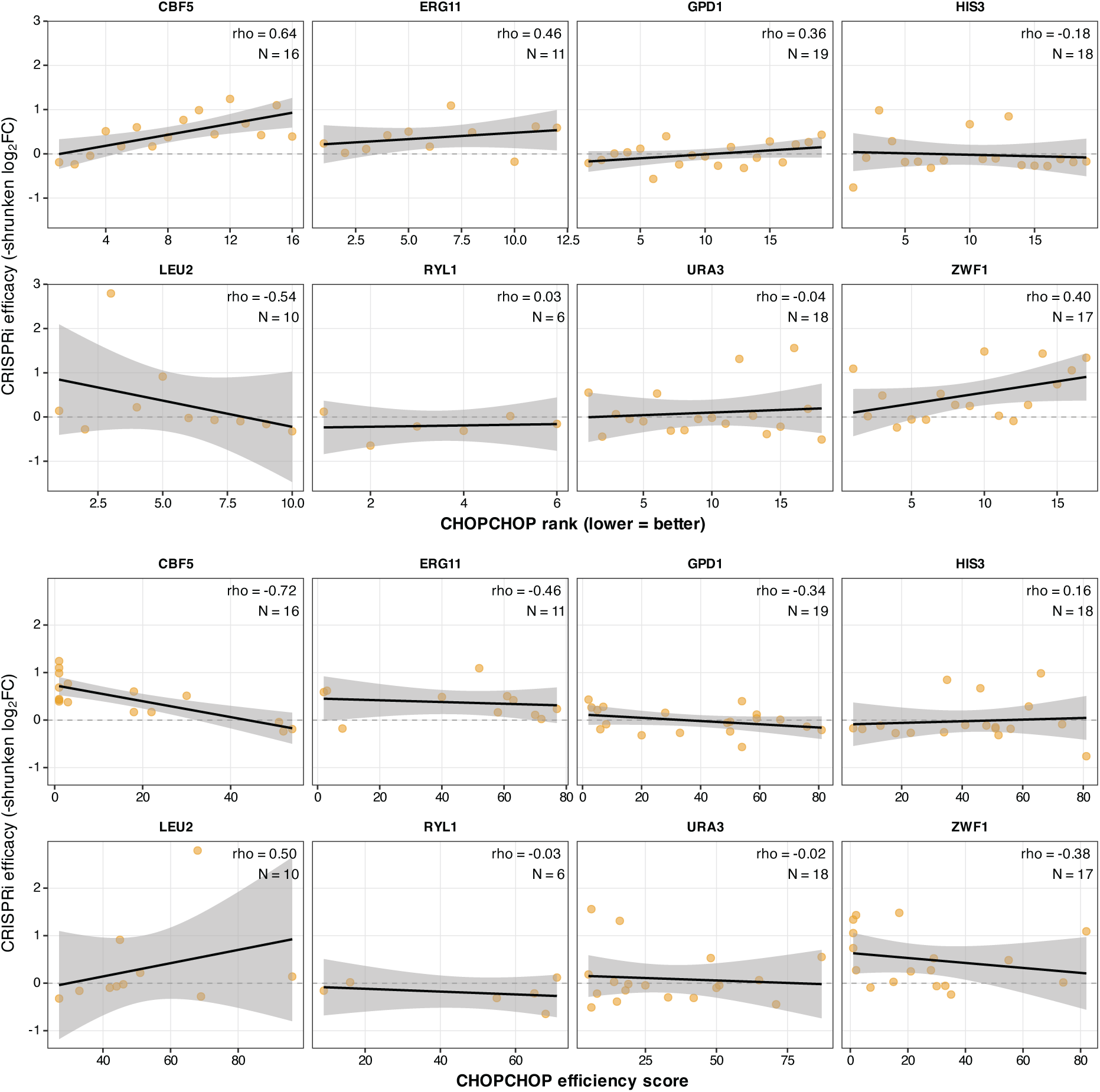
Comparison of CHOPCHOP scores with experimentally measured CRISPRi efficacy. CHOPCHOP **(A)** Rank and **(B)** predicted Efficiency scores are compared with experimentally measured crRNA efficacy (−shrunken log2FC) at G20 in minimal medium. Relationships are shown separately for each target gene. Each point represents an individual crRNA; lines indicate linear fits. Spearman ρ and the number of crRNAs are indicated within each panel.

**Supplementary Figure 9.**
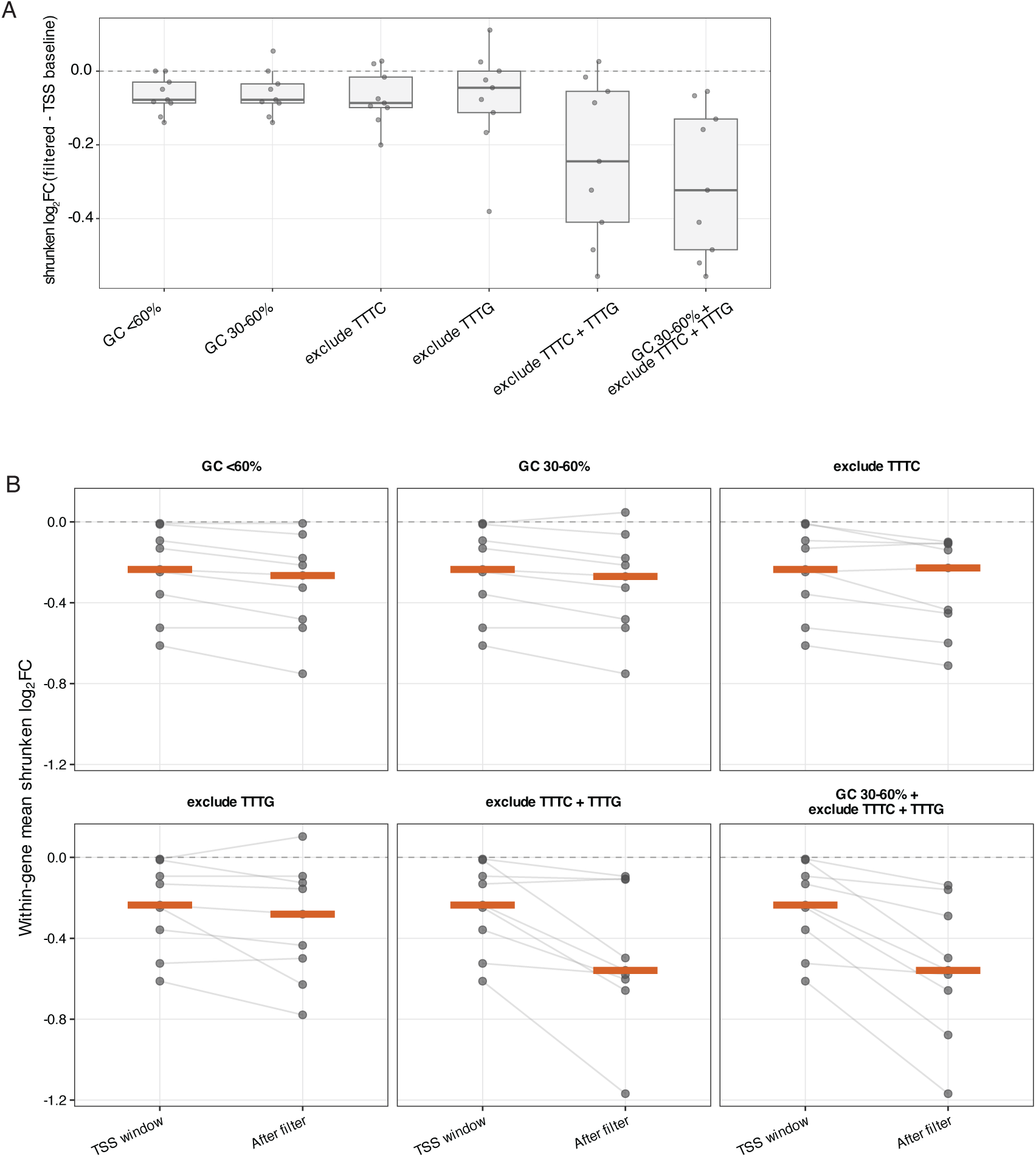
Evaluation of sequence-based crRNA filtering within the preferred TSS-targeting window. **(A)** Change in within-gene mean shrunken log2FC after application of individual and combined GC-content and PAM filters relative to all crRNAs within the −260 to +50 bp TSS window. Each point represents one target gene, and boxplots summarize the distribution across genes. Negative values indicate that filtering enriched for crRNAs showing stronger depletion. **(B)** Within-gene mean shrunken log_2_FC before and after application of the indicated GC-content and PAM filters. The TSS-window value includes all crRNAs within −260 to +50 bp, whereas the post-filter value includes only crRNAs retained by the indicated sequence filter. Lines connect values for the same target gene.

**Supplementary Figure 10.**
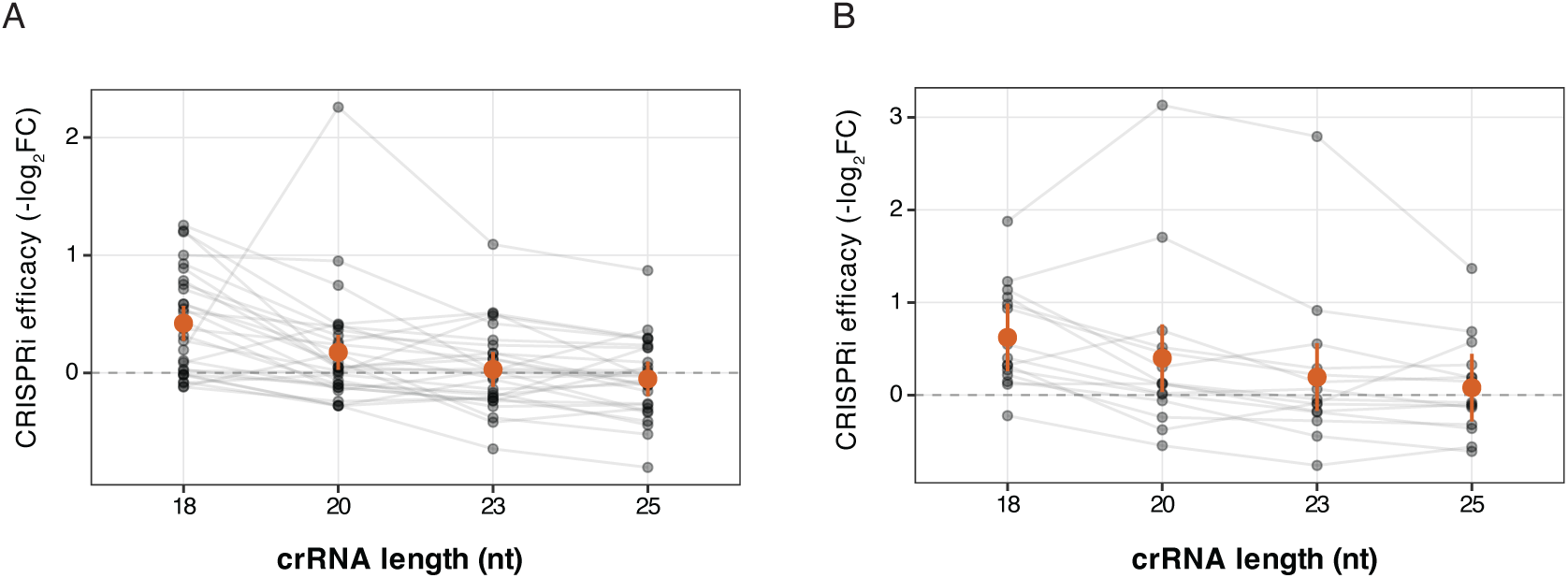
Effect of crRNA spacer length in essential and conditionally essential genes. Spacer-length effects were analyzed separately for **(A)** essential and **(B)** conditionally essential genes using complete matched sets of N18, N20, N23 and N25 crRNAs targeting the same genomic positions. Individual crRNAs are shown as grey points, with lines connecting spacer-length variants targeting the same target site. Orange points and error bars indicate adjusted estimated marginal means and 95% confidence intervals from linear mixed-effects models containing target gene as a fixed effect and target site as a random intercept.

**Table S1.** Target genes in the CRISPRi/a libraries.

| Target gene |  | Effector | Screening conditions | Expected direction for efficient crRNAs |
|---|---|---|---|---|
| ***S. cerevisiae*** **ortholog** | ***Y. lipolytica*** **gene** |  |  |  |
| **Conditionally essential genes** |  |  |  |  |
| *LEU2* | *YALI1_C00464g* | CRISPRi | SCC/SCM | ↘ |
| *HIS3* | *YALI1_E40305g* | CRISPRi | SCC /SCM | ↘ |
| *URA3* | *YALI1_E31685g* | CRISPRi | SCC /SCM | ↘ |
| **Essential genes** |  |  |  |  |
| *CBF5* | *YALI1_E16451g* | CRISPRi | SCC /SCM | ↘ |
| *ZWF1* | *YALI1_E26811g* | CRISPRi | SCC /SCM | ↘ |
| *RYL1* (*SEC4*) | *YALI1_E27207g* | CRISPRi | SCC /SCM | ↘ |
| *ERG11* | *YALI1_B07094g* | CRISPRi | SCC /SCM | ↘ |
| *GUT2 (GPD1)* | YALI1_B18499g | CRISPRi | SCC /SCM | ↘ |
| *ACC1* | *YALI1_C15991g* | CRISPRi | SCC /SCM | ↘ |
| **Drug resistance** |  |  |  |  |
| *CAN1* | *YALI1_B25360g* | CRISPRi | SCC | No change |
|  |  |  | SC + L-canavanine | ↗ |
| *ERG11* | *YALI1_B07094g* | CRISPRa | SCC | No change |
|  |  |  | SC + fluconazole | ↗ |
| **Lipid biosynthesis and accumulation** |  |  |  |  |
| *ACC1* | *YALI1_C15991g* | CRISPRa | Fermentation medium | Enriched in high fluorescence gates |
| *DGA1* | *YALI1_E38810g* | CRISPRa | Fermentation medium | Enriched in high fluorescence gates |

**Table S2.** Growth metrics of dLb-Mxi1 and dHyperLb-Mxi1 strains carrying non-targeting (NT) or *ZWF1*-targeting crRNAs.

| Effector | Target | Metrics |  |
|---|---|---|---|
|  |  | **Mean AUC (OD<sub>600</sub> vs. time) + SD** | **Mean maxOD + SD** |
| dLb-Mxi1 | NT | 12.297 ± 0.460 | 0.796 ± 0.019 |
|  | ZWF1 | 10.272 ± 0.277 | 0.713 ± 0.016 |
| dHyperLb-Mxi1 | NT | 11.736 ± 0.529 | 0.784 ± 0.020 |
|  | ZWF1 | 6.678 ± 0.213 | 0.509 ± 0.025 |

**Table S3.** Plasmids used in this study.

| Name | Description | Parent | Reference |
|---|---|---|---|
| pCfB6576 | Integrative *Y. lipolytica* vector targeting the IntC_2 neutral genomic locus; Amp<sup>R</sup> for selection in *E. coli* and Hyg<sup>R</sup> (hph) for selection in *Y. lipolytica.* Used as the backbone for dCas12a expression plasmids. | - | Addgene #106134 |
| pcfb3405 | Replicative *Y. lipolytica* vector (CEN1/Pot1 promoter + 50bp ORF); Amp<sup>R</sup> for selection in *E. coli* and NAT<sup>R</sup> (clonNAT) for selection in *Y. lipolytica.* Used as the backbone for crRNA expression plasmids. | - | Plasmid #106166) |
| pdenAs | denAsCas12a (*Y. lipolytica* codon-optimized) expression plasmid. | pCfB6576 | This study. |
| pdLb | dLbCas12a (*Y. lipolytica* codon-optimized) expression plasmid. | pCfB6576 | This study. |
| pdenAsVPR | denAsCas12a-VPR (*Y. lipolytica* codon-optimized) expression plasmid. | pCfB6576 | This study. |
| pdenAsMxi1 | denAsCas12a-Mxi1 (*Y. lipolytica* codon-optimized) expression plasmid. | pCfB6576 | This study. |
| pdLbMxi1 | dLbCas12a-Mxi1 (*Y. lipolytica* codon-optimized) expression plasmid. | pCfB6576 | This study. |
| pdLbVPR | dLbCas12a-VPR (*Y. lipolytica* codon-optimized) expression plasmid. | pCfB6576 | This study. |
| pdHyperLbMxi1 | dHyperLbCas12a-Mxi1 (*Y. lipolytica* codon-optimized) expression plasmid. | pCfB6576 | This study. |
| pdHyperLbVPR | dHyperLbCas12a-VPR (*Y. lipolytica* codon-optimized) expression plasmid. | pCfB6576 | This study. |

**Table S4.**
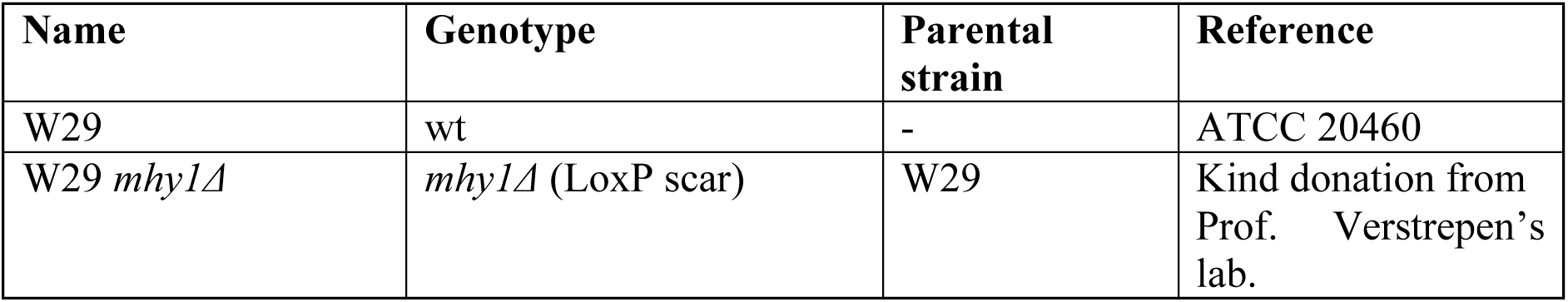
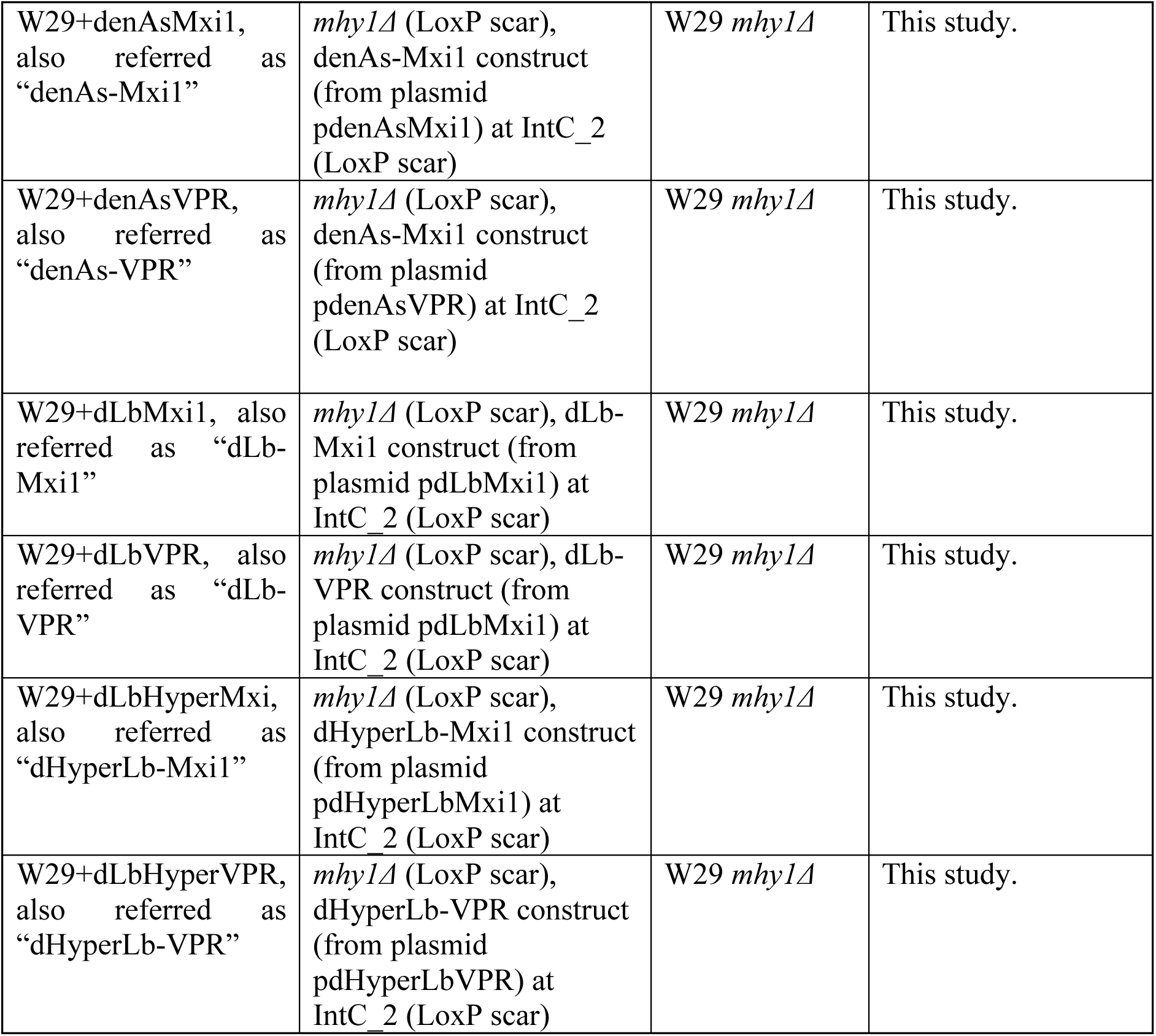
*Y. lipolytica* strains used in this study.

| Name | Genotype | Parental strain | Reference |
|---|---|---|---|
| W29 | wt | - | ATCC 20460 |
| W29 *mhy1Δ* | *mhy1Δ* (LoxP scar) | W29 | Kind donation from Prof. Verstrepen’s lab. |
| W29+denAsMxi1, also referred as “denAs-Mxi1” | *mhy1Δ* (LoxP scar), denAs-Mxi1 construct (from plasmid pdenAsMxi1) at IntC_2 (LoxP scar) | W29 *mhy1Δ* | This study. |
| W29+denAsVPR, also referred as “denAs-VPR” | *mhy1Δ* (LoxP scar), denAs-Mxi1 construct (from plasmid pdenAsVPR) at IntC_2 (LoxP scar) | W29 *mhy1Δ* | This study. |
| W29+dLbMxi1, also referred as “dLb-Mxi1” | *mhy1Δ* (LoxP scar), dLb-Mxi1 construct (from plasmid pdLbMxi1) at IntC_2 (LoxP scar) | W29 *mhy1Δ* | This study. |
| W29+dLbVPR, also referred as “dLb-VPR” | *mhy1Δ* (LoxP scar), dLb-VPR construct (from plasmid pdLbMxi1) at IntC_2 (LoxP scar) | W29 *mhy1Δ* | This study. |
| W29+dLbHyperMxi, also referred as “dHyperLb-Mxi1” | *mhy1Δ* (LoxP scar), dHyperLb-Mxi1 construct (from plasmid pdHyperLbMxi1) at IntC_2 (LoxP scar) | W29 *mhy1Δ* | This study. |
| W29+dLbHyperVPR, also referred as “dHyperLb-VPR” | *mhy1Δ* (LoxP scar), dHyperLb-VPR construct (from plasmid pdHyperLbVPR) at IntC_2 (LoxP scar) | W29 *mhy1Δ* | This study. |

**Table S5.**
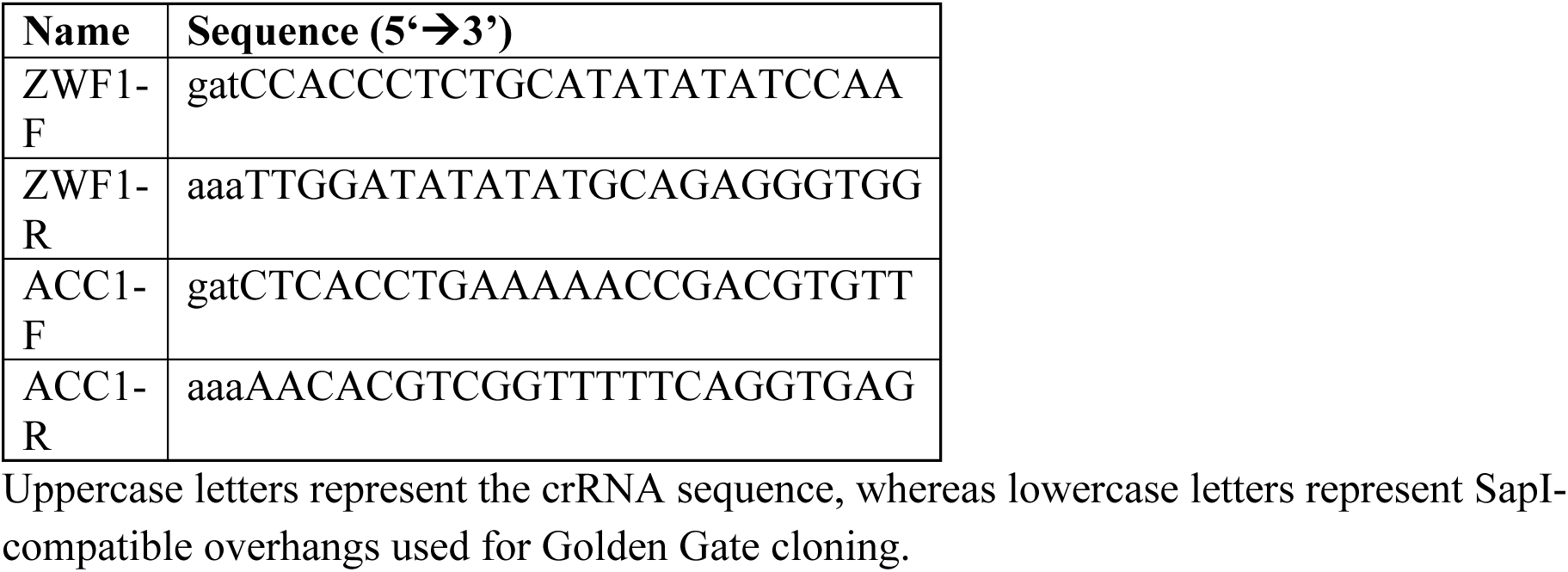
Individual crRNA sequences used in this study.

